# Shifts in embryonic oxygen levels cue heterochrony in limb initiation

**DOI:** 10.1101/2024.10.25.620348

**Authors:** Meng Zhu, ChangHee Lee, Rinaldo Catta-Preta, Clifford J. Tabin

## Abstract

Heterochrony, or the alteration of developmental timing, is an important mechanism of evolutionary change. Avian species display synchronized growth of the forelimbs and hindlimbs, while mammalian species show delayed hindlimb development. We find that mammalian limb heterochrony is evident from the start of limb bud formation, and is associated with heterochronic expression of T-box transcription factors. This heterochronic shift is not due to changes in cis-regulatory sequences controlling T-box gene expression, but unexpectedly, is dependent upon differential oxygen levels to which avian and mammalian embryos are exposed prior to limb initiation. This is mediated, at least partially, by an NFKB transcription factor, *cRel*, and by the oxygen-sensing transcription factor hypoxia inducible factor alpha (*Hif1a*). Together, these results provide mechanistic understanding of an important example of developmental heterochrony and exemplify how the maternal environment regulates timing during embryonic development.

## Introduction

Tetrapod species have two pairs of limbs that are localized at rostral (forelimb) and caudal (hindlimb) positions along the trunk. Although the two pairs of appendages follow the same general developmental blueprint, they display differences in temporal pattern that align with evolutionary lineages ^1^. A particularly interesting case is the amniote clade, where avians and mammals demonstrate disparate temporal dynamics in limb development. In avian species, the early embryonic stages of limb outgrowth are synchronized; while in mammalian species, the hindlimb develops much later than that of the forelimb ^1^. The extent of this heterochrony varies among mammals such that the non-placental mammals display a more pronounced delay than placental mammals, and among the non-placental marsupial species, the ones that have shorter gestation time exhibit higher levels of heterochrony. This heterochronic delay in hindlimb development in mammals has been postulated to represent an "energy trade-off" strategy ^2,3^, which prioritizes the growth of immediately required structures in the context of limited nutrient availability during early developmental stages. However, a molecular-level understanding of limb heterochrony regulation remains entirely unsolved. Indeed, in a more general sense, the control of timing during development is vastly under-studied relative to the control of spatially localized gene expression.

We here investigate the molecular basis of limb heterochrony between avian and mammalian species. We find that limb heterochrony in mammals starts at the limb initiation stage. The delayed onset of hindlimb development can be correlated to expression of *Tbx4*, a transcription factor known to regulate the first steps of hindlimb formation. By coupling RNA-sequencing analyses with gain-of-function screens, we identify a repressive role of the NFKB transcription factor *cRel* in regulating *Tbx4* expression. Unexpectedly, we found that the low oxygen levels that early mammalian embryos experience, a by-product of viviparity, causes upregulation of *cRel*, together with hypoxia inducible factor 1 alpha (*Hif1a*), which attenuate *Tbx4* expression in mammalian hindlimb tissues. Our study presents the first mechanistic understanding of mammalian limb heterochrony, and illustrates how the maternal environment can instruct timing of tissue development in the fetus.

## Results

### Limb heterochrony in mammals starts at the epithelial-to-mesenchymal transition

To interrogate the regulation of limb heterochrony, we first characterized the timing of the earliest steps of limb development in both chick and mouse embryos. Both species initiate limb formation by an epithelial-to-mesenchymal transition (EMT) of cells residing in lateral plate mesoderm (LPM). This leads to the emergence of the paddle-like limb bud, and culminates in the patterning and differentiation of limb progenitor cells into the connective tissues and cartilages. To better understand the stage at which limb heterochrony initially occurs, we first compared the timing of EMT. Our previous work established that the EMT in the chicken embryo takes place in the limb fields at 45 to 52-hour post-gastrulation, or Hamburger Hamilton (HH) stages 12-13 in the forelimb, and 48 to 53-hour post-gastrulation (HH13-14) in the hindlimb ^4^. For the mouse, EMT in the forelimb was previously characterized as occurring between 60-hour post-gastrulation (E8.5) and 66-hour post-gastrulation (E8.75) ^4^. We expanded the analyses to the mouse hindlimb by examining 72-hour post-gastrulation to 96-hour post-gastrulation (E9.0-E10.0) mouse embryos. We found that at 84-hour post-gastrulation (E9.5), majority of the cells in the hindlimb forming regions still remained epithelial, as indicated by polarized F-actin and LAMININ ^4^ and a monolayer organization in the LPM epithelium (Figure 1A). By contrast, at 90-hour post-gastrulation (E9.75), most cells in the hindlimb region showed mesenchymal morphology, with unpolarized distribution of both F-actin and Laminin and the multi-layer stacking of unorganized cells (Figure 1A). These results thus show that EMT initiating the hindlimb in mouse embryos occurs between 84- to 90-hour post-gastrulation (E9.5 and E9.75). Together with previous results, quantifications pointed to a 2-hour difference in EMT timing between forelimb and hindlimb in chicken embryos, in contrast to an 18-hour difference in mice, thus, tracing limb heterochrony in mammals back to the EMT stage of limb development (Figure 1B).

**Figure 1.**
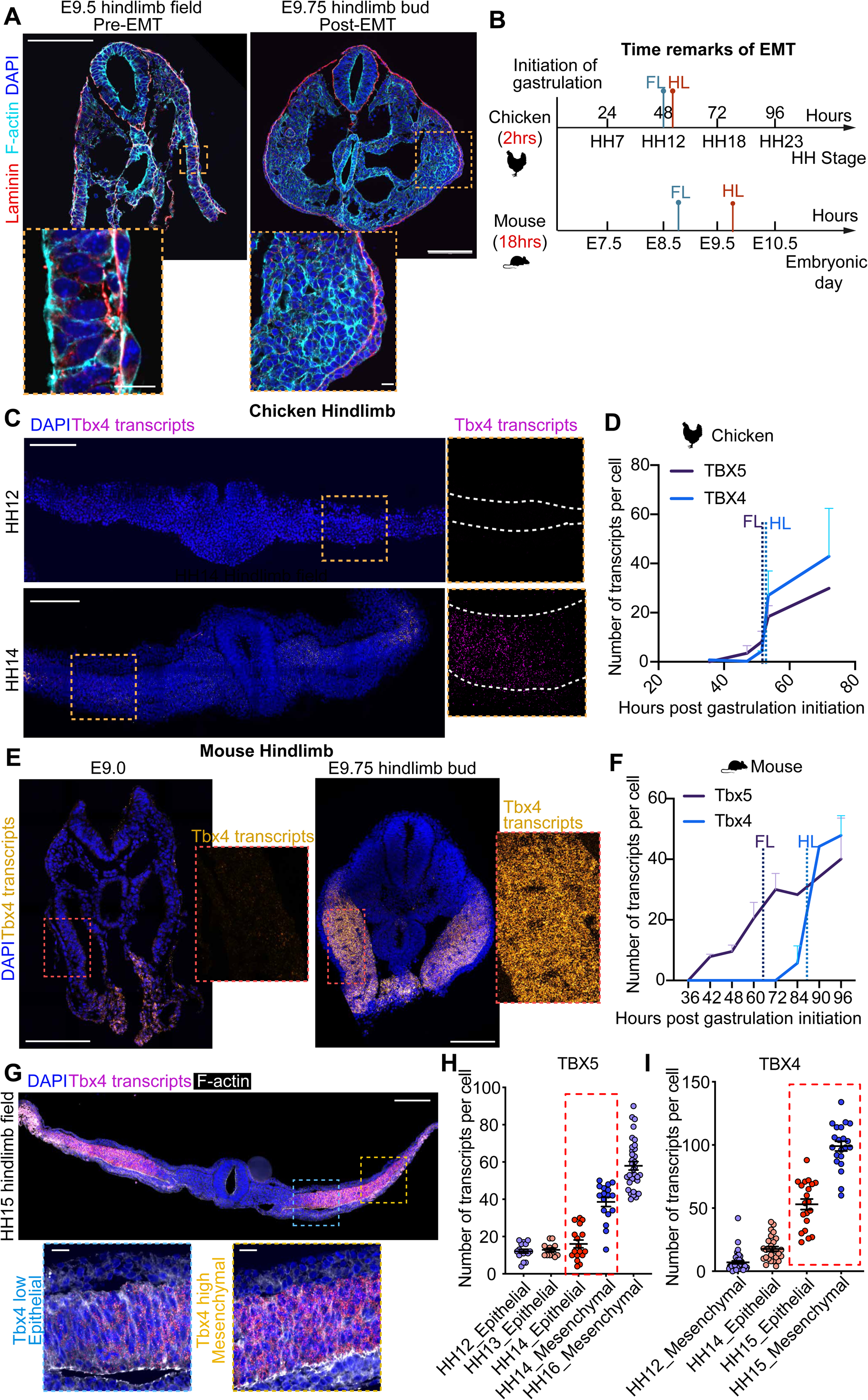
The expression of Tbx5 and Tbx4 correlate with the timing of Epithelial-to-mesenchymal transition (EMT) and the limb heterochrony. (**A**) Representative images of 84-hour post-gastrulation (E9.5) and 90-hour post-gastrulation (E9.75) mouse embryos sectioned at the hindlimb field to stain the F-actin and Laminin DAPI to reveal the epithelial/mesenchymal state of the cells. While at E9.5 the cells remain epithelial shapes, at E9.75, the cell completely transited to mesenchymal state. N=4 embryos for E9.5; N=2 embryos for E9.75. N=2 experiments. (B) Timeline showing the timings of EMT in forelimb and hindlimb and in mouse and chicken embryos. (C) Representative images of chicken hindlimb sections at 47-hour post-gastrulation (HH12) and 51.5-hour post-gastrulation (HH14) revealing the transcripts levels of Tbx4 by Hybridization chain reaction (HCR), and co-stained with DAPI. (D) Quantification of chicken Tbx5 and Tbx4 transcripts levels across 27.5- to 72-hour post-gastrulation (HH8-HH18) stages in forelimb and hindlimb tissues. The stage information is converted to hours post gastrulation initiation at HH4. HH8=27.5h, HH10 =35.5h, HH12=47h, HH14=51.5h, HH16=53.5h and HH18=72h. Dotted lines indicated the EMT timings of forelimb and hindlimb respectively. N=2-6 embryos are used for each tissue and eac h stage. (E) Representative images of E9.0 and E9.75 mouse hindlimb sections revealing the transcripts levels of Tbx4 by Hybridization chain reaction (HCR), and co-stained with DAPI. (F) Quantification of mouse Tbx5 and Tbx4 transcripts levels across E7.5 to E10.0 stages in forelimb and hindlimb tissues. The stage information is converted to hours post gastrulation initiation at HH4. E7.5=36h, E7.75=42h, E8.0=48h, E8.5=60h, E9.0=72h, E9.5=84h, E9.75=90h, E10.0=96h and E10.25=102h. Dotted lines indicated the EMT timing of forelimb and hindlimb respectively. N=2-4 embryos are used for each tissue and each stage. N=2 experiments. (G) Images showing chicken hindlimb at 52.5-hour post-gastrulation (HH15) to stain Tbx4 transcripts by HCR and F-actin and DAPI to correlate with cellular state. Epithelial cells have lower level of Tbx4 than mesenchymal cells. dotted lines indicate magnified areas. (H-I) Quantifications of levels of Tbx5 (H) and Tbx4 (I) in epithelial and mesenchymal cells of chicken forelimb and hindlimb tissues. Dotted lines indicate the transitioning stage when epithelial and mesenchymal cells both present in the same tissue. A clear correlation with the levels of Tbx5 and Tbx4 to epithelial/mesenchymal state can be detected. N=8 embryos, N=2 experiments. Scale bars for overview images, 150μm; scale bars for magnified images, 15μm.

### The heterochrony in EMT correlates with timing of expression of T-box transcription factors

We hypothesized that the discrepancy in timing of EMT between mouse and chicken embryos would be attributable to the heterochronic expression of key limb initiation regulators. To explore this hypothesis, we analyzed the transcriptional dynamics of signaling pathway ligands and transcription factors previously described as regulating limb initiation, including *Fgf10*, *Wnt8c*, *Wnt2b*, *Tbx5* and *Tbx4* ^5–10^. *Fgf10* and *Wnt2b* act downstream of *Tbx5* to initiate forelimb EMT, while *Fgf10* and *Wnt8c* act downstream of *Tbx4* to initiate hindlimb EMT ^6^. To precisely determine their temporal correlation to the timing of EMT, we utilized Hybridization Chain Reaction (HCR) ^11^, a single molecule fluorescence in situ hybridization technology, to visualize transcripts of each gene and to determine their expression dynamics at spatial and temporal resolution. The designed probes show high specificity to the genes of interest as the expression profiles were consistent with results obtained using other methods ^5,12,13^ (Figure 1C,E,G, Figure S1A-D).

Our analysis across developmental stages in the forelimb and hindlimb regions of chicken and mouse embryos revealed that the dynamics of the T-box transcription factors, *Tbx5* and *Tbx4*, exhibited the strongest correlation with the timing of EMT (Figure 1C-F; Figure S1E-G). Specifically, we found that the time when EMT occurs consistently corresponds to the time when half-maximum levels of the *Tbx5* or *Tbx4* transcripts are reached in the limb mesenchyme of both species (Figure 1D,F), a correlation not seen with the other genes tested. Furthermore, we found that this correlation is reflected at the cellular level: cells that transit earlier to the mesenchymal state display higher levels of *Tbx5*/*4* transcripts than those yet to transit (Figure 1G-I; Figure S1E-G). Together, these results suggest T-box transcription factors as the plausible regulators of EMT heterochrony.

Genetic studies show that the depletion of *Tbx5* or *Tbx4* results in forelimb or hindlimb outgrowth defects, while their overexpression can induce ectopic limb growth ^8,9,12,14,15^. To directly test whether *Tbx5*/*4* expression regulates the timing of limb formation, we examined mouse embryos deficient for each of these genes. We had previously shown that depletion of Tbx5 impairs forelimb EMT^4^. We therefore turned our attention to the established Tbx4 knockout mouse line ^9^. At 90-hour post-gastrulation (E9.75), the cells in hindlimb regions of heterozygous *Tbx4* knockout embryos have already undergone EMT, as characterized by the localization of F-actin and Laminin. In contrast, a significant proportion of cells in the *Tbx4* null embryos remained epithelized and localized to the basal region of the LPM (Figure S2A-C). These results, together with the previous studies, show that the limb EMT heterochrony in the mouse embryo is associated with the heterochronic expression of *Tbx5* and *Tbx4*.

### Expression of known upstream activators of *Tbx4* is concordant between chicken and mouse embryos

To investigate the mechanisms leading to delayed *Tbx4* expression in the mouse embryo, we started by examining the dynamics of the signaling pathways and transcription factors known to regulate the activation of *Tbx4* expression. As gastrulation commences, the genes in the Hox clusters are expressed in a 5’ to 3’ order, resulting in the segmented anterior-posterior patterning of the trunk ^16^. The forelimb regions are marked by expression of Hox4 paralogues, while the hindlimb regions are specified by Hox9 paralogues ^17,18^. Interactions with a set of signaling pathways involving Bone Morphogenetic Protein (BMP) activity, Wnt activity, and the inhibition of Retinoic acid (RA) signaling are also required for the specification of posterior LPM ^19,20^. These inputs together activate expression of the transcription factors *Pitx1* and *Isl1* that are enriched in posterior LPM, marking the regions that form the prospective hindlimb ^21–23^. The knockout of each of these transcription factors in the mouse embryo leads to downregulation of *Tbx4* at the hindlimb initiation stage ^24–26^.

We first examined signaling activities essential for LPM specification. Existing literature does not support any notable disparities in the timing of Wnt or RA activation between the mouse and chick ^27–30^. Additionally, our examination of the temporal and spatial distribution of phosphorylated Smad proteins, indicators of BMP pathway activation, revealed no discernible differences between species (Figure 2A-B). Thus, there do not appear to be temporal alterations in signaling pathways required to specify the hindlimb in the two species.

**Figure 2.**
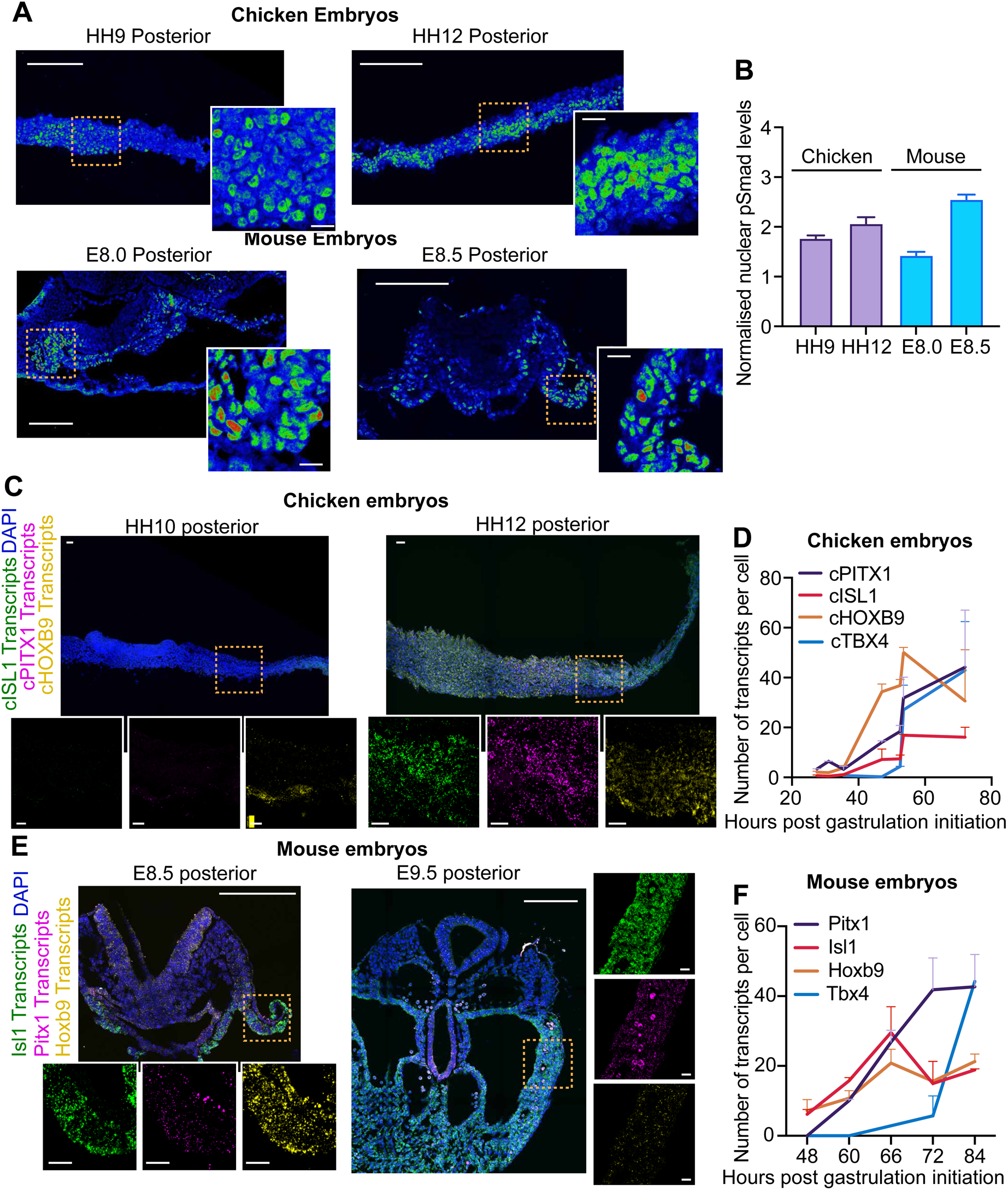
Upstream activators of Tbx4 show concordant expression with Tbx5 between chicken and mouse embryo. (**A**) Representative images of phosphorylated Smad proteins (p-Smad) staining in hindlimb tissues of chicken embryos at HH9, HH12 and mouse embryos at E8.0 and E8.5. Dotted lines indicate the magnified region. (**B**) Quantifications of p-Smad nuclear levels in chicken and mouse hindlimb tissues as in **A**. No significant differences in the kinetics of p-Smad activation are observed between conditions. N=34 cells for HH9 and N=20 cells for HH12 chicken hindlimb tissues. N=2 embryos for each stage. N=60 cells for E8.0 and N=171 cells for E8.5 for mouse embryos. N=2-3 embryos for each stage. N=2 experiments. (**C**) Representative images of PITX1, ISL1 and HOXB9 expression revealed by HCR and co-stained with DAPI in 35.5-hour post-gastrulation (HH10) and 47-hour post-gastrulation (HH12) chicken hindlimb field tissues. Pitx1 and Isl1 are robustly expressed from 47-hour post-gastrulation (HH12) chicken embryo. Dotted lines indicate magnified regions. (**D**) Quantifications of chicken PITX1, ISL1 and HOXB9 transcripts levels by HCR across HH8-18 stages of hindlimb tissues as in **a**. 2-4 embryos for each stage quantified. N=2 experiments. (**E**) Representative images of Pitx1, Isl1 and Hoxb9 expression revealed by HCR and co-stained with DAPI in E8.5 and E9.5 mouse hindlimb field tissues. Pitx1 and Isl1 are robustly expressed from E8.5 in mouse embryo. Dotted lines indicate magnified regions. (**F**) Quantifications of mouse Pitx1, Isl1 and Hoxb9 transcripts levels by HCR across E7.5-E10.0 stages of hindlimb tissues as in **C**. 2-3 embryos for each stage, N=2 experiments. For **B** and **D**, data are shown as mean ± S.E.M. Data is shown as mean ± S.E.M. Scale bars for overview images, 150μm; scale bars for magnified images, 15μm.

We then probed the transcriptional dynamics of *Hoxb9*, *Pitx1* and *Isl1* expression in mouse and chicken embryos by HCR (Figure 2C-F), in terms of hours post-gastrulation, and used the time to reach half-maximum as a metric for the timing of gene expression. We found that *Hoxb9*, *Pitx1*, and *Isl1* activation times were similar in both species: in the chick, where the forelimb and hindlimb initiate at nearly the same time, expression of *Tbx5*, *Tbx4*, *Pitx1* and *Isl1* are all initiated simultaneously. Conversely in the mouse, where hindlimb initiation is delayed relative to the forelimb, *Hoxb9*, *Pitx1* and *Isl1* are still activated together with *Tbx5,* despite the delay of Tbx4 (Figure S2D-E; Figure 2D,F). These data suggest that the delay in Tbx4 expression in the mouse embryo (22-hour after *Hoxb9*/*Isl1*/*Pitx1* in mouse vs 2-hour in chicken) is not a consequence of delayed specification of the hindlimb lateral plate mesoderm (LPM), or a lag in the expression of known upstream activators.

### Shifted timing of Tbx4 activation is not due to alterations in associated cis-regulatory elements

Heterochrony is an important mode of evolutionary change. The modern model posits that evolutionary change is largely attributable to mutations in cis-regulatory elements (CREs) ^31^. To determine whether CRE changes proximal to *Tbx4* are responsible for the shifted timing of that gene’s expression in the mouse, we sought to compare *Tbx4* enhancer activity between mouse and chicken. For the mouse embryo, previous studies identified two genomic regions, named HLEA and HLEB, as necessary and sufficient to activate *Tbx4* expression in the developing hindlimb ^32^. To identify chicken Tbx4 enhancers, we took advantage of an ATAC-seq analysis previously carried out in the lab ^33^, and selected regions that showed accessibility in 72-hour post-gastrulation (HH18) hindlimb mesenchyme but not in the forelimb or flank. This criterion filtered the pool into 4 candidate genomic regions located within 100kb of *Tbx4* transcriptional start site (Figure 3A, Figure S3A). To test whether these regions can drive the transcription in a pattern resembling that of endogenous *Tbx4*, we electroporated enhancer:Citrine reporter constructs, each carrying a candidate region, into chicken forelimb, flank or hindlimb tissues at 51.5-hour post-gastrulation (HH14) (Figure 3B). When the electroporated embryos were examined at 72-hour post-gastrulation (HH18), 1 of the 4 candidate regions drove strong signal, restricted to the hindlimb tissue (Figure 3C-D; Figure S3A). This enhancer was designated as Tbx4-Rec1. To reveal the temporal activation pattern of Tbx4-Rec1, we quantified Citrine signal in hindlimb tissues of different stages by flow cytometry. This revealed that Tbx4-Rec1 activity is initiated at 47-hour post-gastrulation (HH12) and becomes strong by 53.5-hour post-gastrulation (HH16), paralleling the timing of endogenous *Tbx4* expression (Figure 3E; Figure S3B-E). Thus, Tbx4-Rec1 activity resembles the expression profile of *Tbx4* both spatially and temporally in the chicken embryo (Figure 1C-D). Finally, to determine whether Tbx4-Rec1 is functionally required for Tbx4 expression, we inhibited the endogenous Tbx4-Rec1 region by deploying the dCas9-KRAB assay, which utilizes a catalytically dead CAS9 fused to the Krüppel-associated box (KRAB) transcriptional repression domain. This system, guided by specifically designed gRNAs targeting Tbx4-Rec1, acts to methylate the genomic region, thereby repressing it ^34^. We electroporated a construct carrying a gRNA targeting the middle position of Tbx4-Rec1 and the coding sequence of dCas9-krab, into the 47-hour post-gastrulation (HH12) hindlimb field of chicken embryos. After 12 hours, the electroporated cells showed significant reduction of Tbx4 transcripts when compared to non-electroporated counterparts, or cells electroporated with a control gRNA (Figure 3F-I). Together, these experiments implicate Tbx4-Rec1 as a critical enhancer for Tbx4 expression in the embryonic chicken hindlimb.

**Figure 3.**
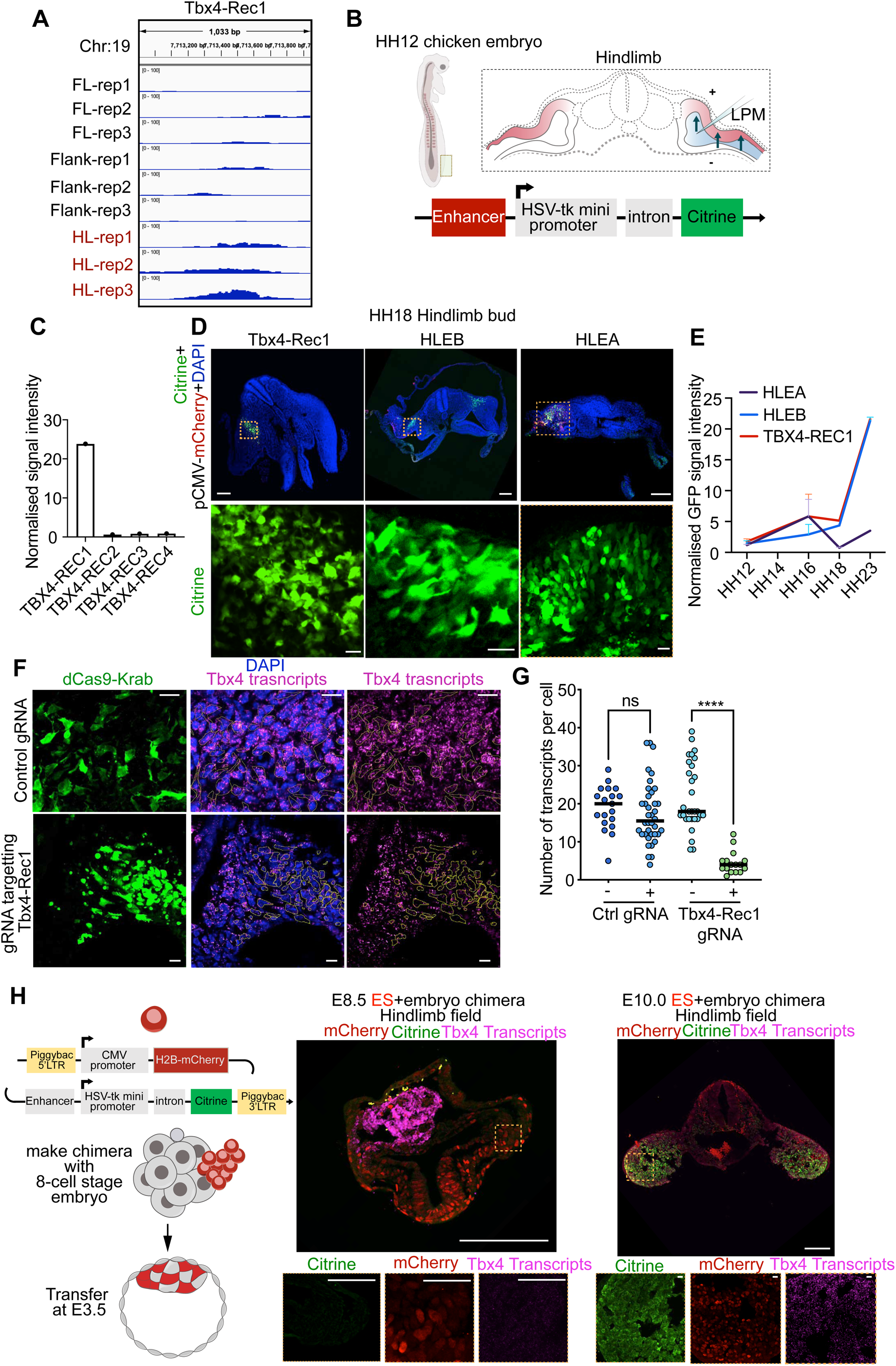
cis-regulatory elements responsible for Tbx4 activation are conserved between chicken and mouse embryos. (**A**) Chromatin accessibility of genomic regions of chicken Tbx4 enhancer designated as Tbx4-Rec1. Data is extracted from ATAC-seq experiments in Young et al., 2019^33^. Tbx4-Rec1 genomic region is higher accessible in hindlimb tissues than forelimb and flank tissues in 72-hour post-gastrulation (HH18) chicken embryos. (**B**) Scheme of enhancer bashing assay. Enhancer:reporter constructs carrying each enhancer candidates for chicken Tbx4 expression were electroplated in 47-hour post-gastrulation (HH12) chicken hindlimb fields. pCMV-mCherry construct is co-electroplated as electroporation control. The enhancer activity was examined at 72-hour post-gastrulation (HH18) and 96-hour post-gastrulation (HH23). (**C**) Quantification of the enhancer activity of the 4 chicken Tbx4 enhancer candidates at 72-hour post-gastrulation (HH18) by flow cytometry. Enhancer activity is calculated as the signal ratio of Green channel/Red channel. Data is shown as mean value for each condition. N=10 embryos disassembled for each group, N=3 experiments. Only Tbx4-Rec1 show high activation. (**D**) Representative images of chicken hindlimb tissues electroplated with Tbx4-Rec1:Citrine constructs and sectioned at 72-hour post-gastrulation (HH18) and 96-hour post-gastrulation (HH23) to examine the enhancer activity by Citrine and DAPI. (**E**) Quantifications of mouse and chicken Tbx4 enhancer activities by flow cytometry of chicken hindlimb cells electroplated by each of the enhancer. Data shown as mean± S.E.M N=30 embryos were disassembled for each condition in total, N=3 experiments. (**F**) Representative images of 53.5-hour post-gastrulation (HH16) chicken hindlimb cells electroplated by dCas9-Krab constructs containing control gRNA or with gRNA targeting Tbx4-Rec1 for 13h, and fixed and visualized Tbx4 transcripts by HCR and co-stained with ZsGreen and DAPI. Yellow circles delineate electroplated cells. (**G**) Quantifications of Tbx4 transcripts in conditions shown in **F**. Tbx4 transcripts are significantly downregulated specifically in cells transfected with Tbx4-Rec1 gRNA. ****p<0.0001, ns, not significant, Mann-Whitney test. Data shown as individual points with mean± S.E.M. Each dot indicates one cell. N=19 cells for Ctrl gRNA Citrine-; N=38 cells for Ctrl gRNA Citrine+; N=30 cells for Tbx4-Rec1 gRNA Citrine-; N=16 cells for Tbx4-Rec1 gRNA Citrine+. N=2-3 embryos for each condition, N=2 experiments. (**H**) Scheme for the strategies for producing ES cell-embryo chimera. Wildtype ES cells were transfected with constructs containing piggybac transposase and piggybac LTRs enclosing enhancer:Citrine cassette and CMV:H2B-mCherry cassette. The transgenic ES cells were then aggregated with E2.5 (8-cell stage) mouse embryos and cultured until E3.5. The chimeric embryos were then transferred to pseudo-pregnant females and dissected at deserved stages before and after Tbx4 expression. The representative images of ES-embryo chimeric embryos carrying Tbx4-Rec1:Citrine transgenic ES cells recovered at E8.5 and E10.0 to reveal the enhancer activity by Citrine and mCherry, and co-visualized Tbx4 transcripts by HCR and DAPI. N=6 embryos. N=2 experiments. Scale bars for overview images, 150μm; scale bars for magnified images, 15μm.

Sequence analyses suggest that the 3’ DNA sequence of Tbx4-Rec1 highly resembles that of the mouse Tbx4 enhancer, HLEB (Figure S3F-H). This suggests that the chicken Tbx4-Rec1 and mouse HBEB Tbx4 enhancers are homologous sequences. There is no obvious counterpart to HLEA in the chick genomic sequence near the Tbx4 gene ^32^. To test whether the mouse and chick enhancers are functionally conserved, we performed enhancer-swap experiments to orthogonally express the Tbx4 enhancers across species. To introduce the mouse Tbx4 enhancer into the chicken embryo, we performed electroporation experiments with Citrine reporter constructs carrying either HLEA or HLEB, as above (Figure 3D-E). We found that both HLEA and HLEB were strongly activated, driving reporter expression at a similar level to Tbx4-Rec1 (Figure 3D). Moreover, the timing of activation of both mouse enhancers was similar to that of Tbx4-Rec1 (Figure 3E, Figure 1D). To introduce the chicken Tbx4 enhancer into the mouse embryo, we produced Embryonic Stem cells (ES)-embryo chimeras (Figure 3H). We first generated transgenic mouse ES cell lines each with a random genomic insertion of a Tbx4-Rec1:Citrine reporter cassette (Figure 3H). The transgenic ES cells were then aggregated to 8-cell stage mouse embryos, which were transferred to pseudo-pregnant female mice, and recovered at stages before and after *Tbx4* expression. We optimized the ES cells aggregation protocol to achieve nearly full donor chimerism (Figure S3G, *Methods*). We first examined HLEB:Citrine activity as a positive control. As expected, no Citrine signal can be detected in the hindlimb LPM region from embryos recovered at 66-hour post-gastrulation (E8.75) (Figure S3H); However, strong activity can be detected only in hindlimb but not forelimb regions at 90-hour post-gastrulation (E9.75) (Figure S3I). Such an activation pattern recapitulates the endogenous mouse Tbx4 expression. We then examined Tbx4-Rec1:Citrine activity, and found that, similar to HLEB:Citrine, no Citrine signal was seen at 60-hour post-gastrulation (E8.5), but strong activation was detected at 96-hour post-gastrulation (E10.0) in the hindlimb (Figure 3H). The inter-species enhancer analysis experiments thus demonstrate that the mouse and chicken *Tbx4* enhancers are functionally equivalent, and that any sequence variation between them are insufficient to account for the temporal shift in *Tbx4* expression observed in the mouse embryo.

### NFKB transcription factors are negative regulators of Tbx4 expression

Having ruled out the contribution of cis-regulatory elements to the delayed expression of *Tbx4* in the mouse, we turned into the alternative model that differentially expressed trans-acting factors are responsible for difference in the timing of *Tbx4* expression between the two species. To address this hypothesis, we conducted bulk RNA-sequencing of forelimb and hindlimb tissues from 60-hour post-gastrulation (E8.5) and 90-hour post-gastrulation (E9.75) mouse embryos, as well as forelimb and hindlimb tissues of 47-hour post-gastrulation (HH12) chicken embryos (Figure S4A).

Consistent with literature, our sequencing analyses identified *Tbx5* and *Raldh1a2* as significantly enriched in forelimb tissues ^12,35^, and *Cdx2*, *Evx1*, *Pitx1* as enriched in the hindlimb tissues across both species ^25,36,37^ (Figure S4A). Comparing between stages, we found about 200 genes differentially expressed between 60-hour post-gastrulation (E8.5) and 90-hour post-gastrulation (E9.75) hindlimb tissues in the mouse.

To identify differentially expressed genes that might be functionally important for *Tbx4* expression, we reviewed the documented function of each gene, and filtered out those with known discordant expression in HH12 chicken hindlimb, narrowing the focus to 22 factors related to metabolism, paracrine signaling, and transcriptional regulation (Table S1). To test whether the expression of any of these factors impact *Tbx4* expression, we mis-expressed each gene in the chicken hindlimb. We designed two types of assays based on the gene’s expression profile: 1) potential inhibitors: genes with higher expression at 60-hour post-gastrulation (E8.5) than 84-hour post-gastrulation (E9.5) were overexpressed in the HH12 chicken hindlimb field, and their impact on *Tbx4* expression assessed at 96-hour post-gastrulation (HH23). 2) potential activators: genes upregulated from E8.5 to 90-hour post-gastrulation (E9.75) were overexpressed in gastrulating HH4 chicken embryos to evaluate potential advancement in Tbx4 expression by 35.5-hour post-gastrulation (HH10).

This screen identified *cRel* as a potent repressor of *Tbx4* expression (Figure 4D-F; Figure S4B-C). *cRel* is an effector of the NFKB signaling pathway involved in immune and stress response. We find that overexpression of *cRel*, but not the other candidates that we tested, results in a marked reduction in hindlimb bud size and consistently lower *Tbx4* transcript levels only in electroporated cells, suggesting a cell-autonomous effect of REL in blocking *Tbx4* expression (Figure 4D; Figure S4B-C).

**Figure 4.**
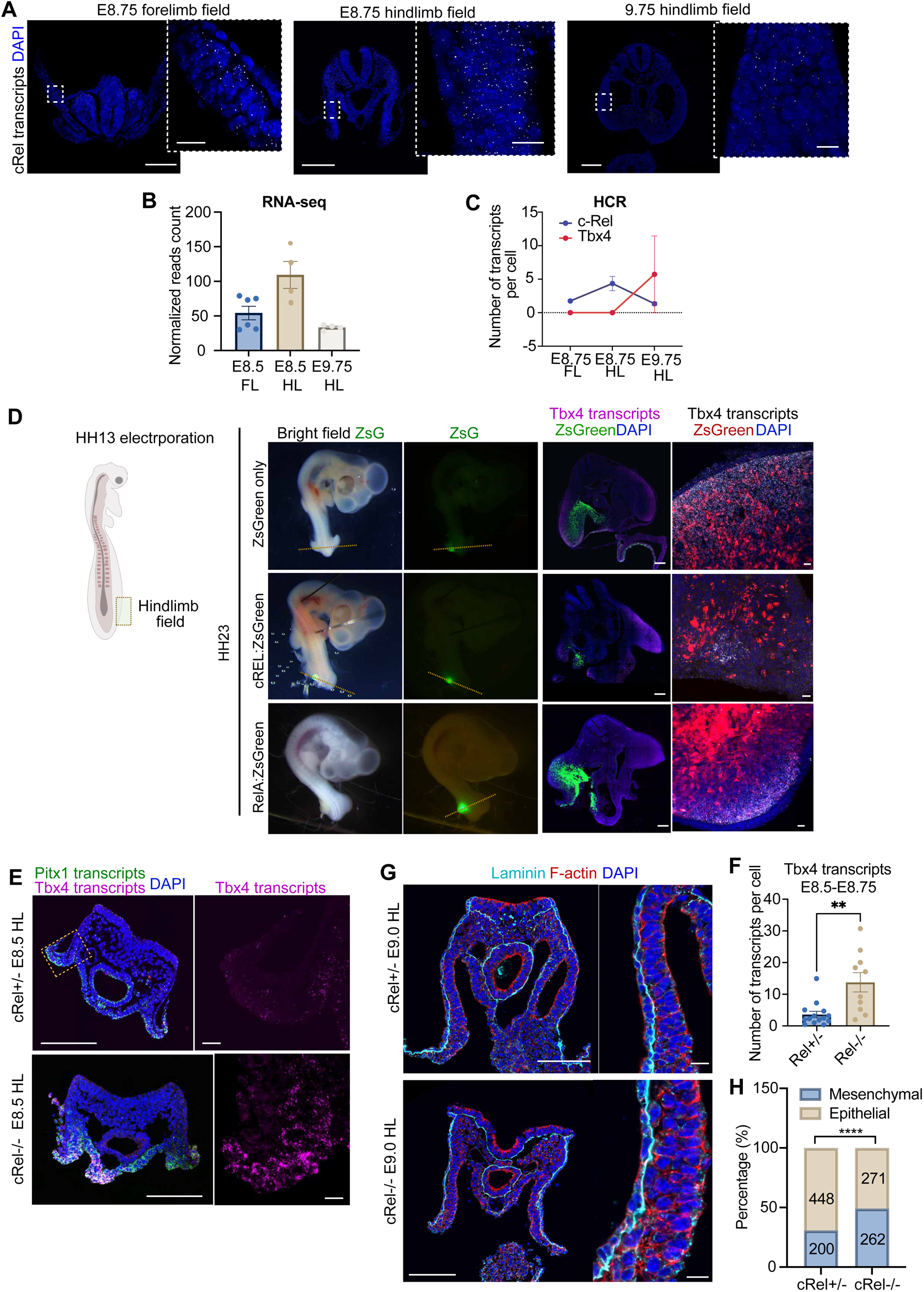
Higher cRel expression represses Tbx4 expression in the hindlimb of mouse embryo. (**A**) Representative images of E8.75 and E9.75 mouse embryos visualizing cRel expression by HCR (co-stained with DAPI) in forelimb and hindlimb tissues. N=2 embryos for each condition. N=2 experiments. (**B**-**C**) RNA-sequencing data (**B**) and HCR quantification (**C**) of cRel expression in E8.5 forelimb and hindlimb tissues and E9.75 hindlimb tissues. cRel expression is downregulated from E8.5 to E9.75 hindlimb tissue. For **b**, N=10 embryos were pooled for each repeat, tissue and stage. N=4 repeats. For **c**, N=2 embryos for each condition. N=2 experiments. (**D**) Representative images of chicken embryos electroplated with ZsGreen only (as a control), cRel:ZsGreen or RelA:ZsGreen into HH13 hindlimb tissue of chicken embryos and cultured for 2 days (at HH23) to examine the hindlimb bud formation and Tbx4 expression by HCR (and co-stained with DAPI). cRel or RelA overexpression significantly reduced the hindlimb bud formation only in the electroplated side of the embryo, and resulted in significant downregulation of Tbx4 expression only in the electroporated cells. N=8 embryos for ZsGreen, N=13 embryos for cRel:ZsGreen, and N=6 embryos for RelA. N=3 experiments. Dotted lines indicate locations of the cross-sections. (**E**) Representative images of the posterior section of cRel heterozygous and homozygous knockout embryos dissected at E8.5, visualized Tbx4 by HCR, and co-stained with DAPI. Pitx1 transcripts were visualized by HCR to reveal the location of the hindlimb field region. Dotted lines indicate magnified regions. (**F**) Quantifications of Tbx4 transcripts in the hindlimb field of conditions shown in **E**. **p<0.01, Mann Whitney test. Each dot represents an embryo. Data is shown as mean ± S.E.M. N=7 experiments. (**G**) Representative images of cRel heterozygous and homozygous knockout embryos dissected at E9.0 and examined the epithelial or mesenchymal state of the hindlimb field cells by the immunostaining of F-actin, Laminin and DAPI. Dotted lines indicate magnified regions. (**H**) Quantifications of epithelial/mesenchymal cells in hindlimb tissues from conditions in **G**. ****p<0.0001, Fisher’s exact test. Data is shown as bar chart, numbers in each bar represents the number of cells examined. N=7 embryos for cRel knockout, N=7 embryos for cRel heterozygous mutants. N=2 experiments. Scale bars for overview images, 150μm; scale bars for magnified images, 15μm.

Analysis of *cRel* knockout mouse embryos from 60-hour post-gastrulation (E8.5) to 84-hour post-gastrulation (E9.5) revealed that *cRel*-null embryos exhibited significant elevation of *Tbx4* expression in hindlimb tissues at E8.5 compared to heterozygous littermates (Figure 4E-F). However, this ectopic expression of *Tbx4* was not observed across all cells, and predominantly occurred on only one side of the embryo (Figure 4E). Despite this heterogeneous expression, the increased *Tbx4* levels were sufficient to advance EMT timing by 72-hour post-gastrulation (E9.0), as *cRel* null hindlimb tissues show a significant higher proportion of mesenchymal cells than heterozygous at E9.0 (Figure 4G-H). Possibility due to the partial penetration, we found no difference in hindlimb bud size between *cRel* null and heterozygous littermates at 84-hour post gastrulation (E9.5) (Figure S4D-E).

One possible explanation for the partial penetrance we observed could be redundancy with related factors co-expressed in the limb bud. Besides *cRel*, the NFKB family includes 4 other transcription factors, *Rela*, *Relb*, *Nfkb1* and *Nfkb2*. Of the 5 factors, *cRel*, *Rela* and *Nfkb1* were detectable in 60-hour post-gastrulation (E8.5) mouse hindlimb tissues in bulk RNA-seq (Figure S4B). To test whether other NFKB transcription factors similarly inhibit Tbx4 expression, we overexpressed *Rela* or *Nfkb1* in chicken hindlimb. We found that the overexpression of *Rela*, but not *Nfkb1*, led to significant reduction of hindlimb bud size (Figure 4D). Examination of *Tbx4* expression revealed a significant downregulation of Tbx4 transcripts in cells overexpressing *Rela*. By analyzing a recently published single-cell RNA sequencing data set, we found that the expression dynamics of *cRel* and *Rela* were similarly downregulated from 60-hour post-gastrulation (E8.5) to 84-hour post-gastrulation (E9.5) in the LPM ^38^. These data suggest *Rela* as a potential compensatory factor that may delay *Tbx4* expression akin to *cRel*.

Altogether, the top-down approach using RNA-sequencing analyses reveals a previously unreported inhibitory role of NFKB transcription factors in delaying *Tbx4* expression in the developing mouse hindlimb bud.

### Maternal hypoxia impedes Tbx4 expression in the hindlimb

Building on our findings that the downregulation of *cRel* and *Rela* from pre- to post-hindlimb initiation stages may be responsible for the delayed activation of *Tbx4* in mice, we next sought to elucidate the regulatory mechanisms governing their temporal expression. One parameter that caught our attention as potentially being different in the two settings was the level of oxygen present. Hypoxia can induce the expression of NFKB transcription factors in a number of contexts ^39,40^, and importantly, evidence suggest that chicken eggs develop in environment close to normoxia (20% oxygen) whereas early mouse embryos are in hypoxia (2%-5% oxygen) ^41–43^. Notably, there is a transition in the mouse embryo from low oxygen levels to normoxia as the placental connection to the maternal circulation becomes robust, at a time correlating with hindlimb initiation. Correspondingly, while chicken embryos exhibit low levels of nuclear HIF1A in the hindlimb field at HH12, mouse embryos at the comparable stage, 60-hour post-gastrulation (E8.5), showed high levels of nuclear Hif1a. Nuclear HIF1A is diminished one day later, at 84-hour post-gastrulation (E9.5), coinciding with robust Tbx4 expression and hindlimb EMT initiation (Figure 5A-C).

**Figure 5.**
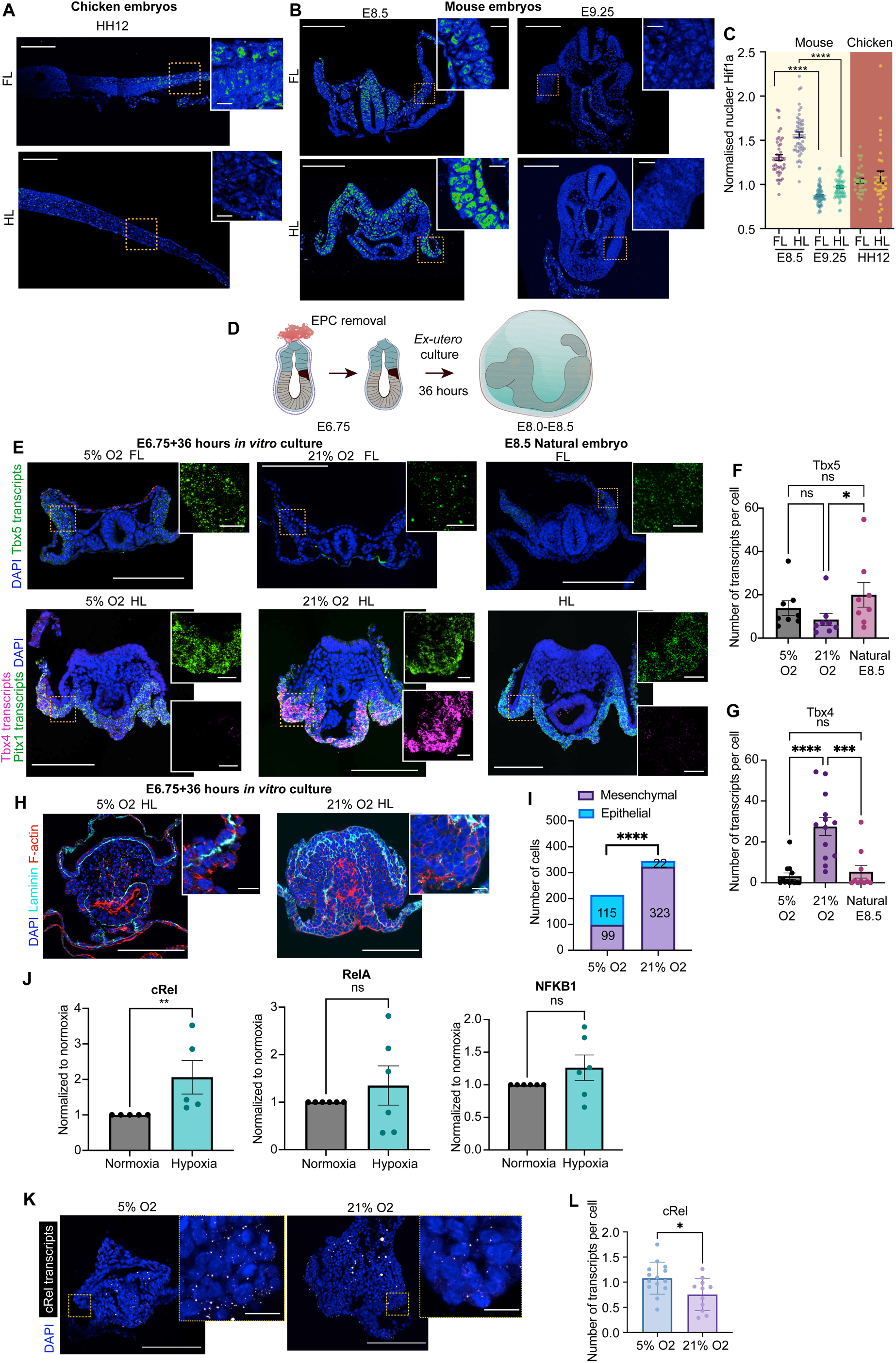
Level of oxygens regulate the timing of Tbx4 expression partially through NFKB transcription factors’ expression. (**A**-**B**) Representative images of Hif1a staining in forelimb and hindlimb tissues of 47-hour post-gastrulation (HH12) chicken embryo (**A**) and E8.5 and E9.25 mouse embryos (**B**). Dotted lines indicate magnified regions. (**C**) Quantifications of the levels of nuclear Hif1a (normalized to DAPI) in conditions shown in a-b. ****p<0.0001, One way ANOVA test. Data shown as mean ± S.E.M, each dot indicates a cell. N=2 embryos for each stage. N=2 experiments. (**D**) Scheme of *ex-utero* culture system. E6.75 mouse embryos were recovered and the ecto-placental cone was removed, the embryos were then cultured in 5% or 20% O_2_ for 1.5 days (36h), after which the embryos reach the stage the morphology of which assemblies that of E8.5 mouse embryo. (**E**) *Ex-utero* cultured embryos in 5% or 20% O_2_ level as in (**D**) were fixed to examine transcripts levels of Tbx5, Pitx1 and Tbx4 (co-stained with DAPI) in forelimb and hindlimb regions after 1.5 days of culture. Natural E8.5 embryos were compared as control. Dotted lines indicate magnified regions. (**F**-**G**) Quantifications of Tbx5 (**F**) and Tbx4 (**G**) in forelimb and hindlimb tissues from conditions shown in **E**. ns, not significant, *p<0.05, ***p<0.001, ****p<0.0001, One way ANOVA test. Data shown as mean ± S.E.M. Each dot indicates an embryo. N=8 experiments for both 5% and 20% O_2_. (**H**) Representative images of *ex-utero* cultured embryos in 5% and 20% O_2_ for 2 days and immunostained with laminin, F-actin and DAPI for revealing epithelial/mesenchymal state of the hindlimb cells. (**I**) Quantification of number of epithelial and mesenchymal cells for conditions shown in **H**. ****p<0.0001, Fisher’s exact test. Data is shown as bar chart, numbers in each bar represents the number of cells examined. N=4 embryos for 5% O_2_; N=3 embryos for 20% O_2_. N=2 experiments. Significant higher proportion mesenchymal cellsare seen in hindlimb cells cultured in 20% O_2_. (**J**) Quantitative PCR (qPCR) analysis of cRel, RelA, NFKB1 expression in embryos ex-utero cultured in 5% or 20% O_2_. **p<0.01, ns, not significant. Mann-Whitney test. Data shown as mean ± S.E.M. Each data point represents one experiment. Hypoxia culture induces significant upregulation of *cRel*. (**K**) *Ex-utero* cultured embryos from E6.5 for over 48 hours in normoxia or hypoxia conditions, processed with HCR targeting cRel transcripts and stained with DAPI. (**L**) Quantification of *cRel* transcripts in conditions shown in **K**. ****p<0.0001, Mann-Whitney test. Data shown as mean ± S.E.M. Each data point represents one embryo. N=5 experiments. Scale bars for overview images, 150μm; scale bars for magnified images, 15μm.

To test whether oxygen levels could play a role in regulating the timing of *Tbx4* expression in mouse embryos, we optimized an *ex-utero* culture system that allowed gastrulating embryos, dissected at 12-hour post-gastrulation (E6.5), to develop until approximately 66-hour post-gastrulation (E8.75) (Figure 5D; Figure S5A; Methods). The culture method is based on previous protocols ^42,44^ but with two modifications: 1) 5% FBS was added into the culture medium as an alternative to human cord serum; 2) the ecto-placental cone (EPC) was removed to enhance oxygen equilibration within and surrounding the embryonic tissues. The latter modification was based on the observation that in the presence of an intact EPC, *Hif1a* expression was not completely extinguished in embryos cultured in normoxic conditions (Figure S5B), likely due to inefficient gas penetration of the yolk sac layer. The cultured embryos generally attain a similar morphology to naturally developed E8.5 embryos in the formation of head-fold, allantois, and a clear distinction of anterior and posterior regions. There were occasional defects in the epithelization of somite tissues, which is likely due to the lack of sufficient mechanical cues in culture ^45^. Nonetheless, we found robust expression of forelimb field and hindlimb field markers *Tbx5* and *Pitx1*, indicating normal differentiation of LPM lineages (Figure 5E; Figure S5C-D; Figure S6A-D).

Leveraging the *ex-utero* culture system, we compared gene expression under different oxygen conditions. We recovered embryos at E6.75, and cultured them for 1.5 days, in either 5% or 20% oxygen, which correspond, respectively, to hypoxic and normoxic conditions. Remarkably, we found consistent activation of *Tbx4* transcripts in mouse embryos cultured in normoxic conditions, which did not occur in those cultured in hypoxic conditions (Figure 5E,G, Figure S6D). Moreover, this activation of *Tbx4* in normoxia-grown embryos is more pronounced than seen in *cRel* knockout embryos. Nearly all these embryos showed premature, elevated, bilateral expression of *Tbx4* (Figure 5E-G). The normoxia cultured embryos additionally show significant upregulation of *Pitx1* in the posterior LPM (Figure S5C; Figure S6D). This effect was specific to these hindlimb genes as, in contrast, the normoxia cultured embryos showed no significant change in *Tbx5* expression (if anything, there was a trend towards down-regulation) (Figure 5F).

Consistent with the ectopic activation of *Tbx4*, we found that the *Tbx4* enhancer Tbx4-Rec1, also showed scattered, yet significant upregulation in the hindlimb field of embryos cultured in normoxia, relative to those grown in hypoxia (Figure S5E-F). Together, these results suggest that forced elevation of oxygen levels result in precocious hindlimb induction, including *Tbx4* expression and EMT, in mouse embryos. We could not carry out the reciprocal experiment, and ask whether culture in hypoxia delays hindlimb initiation in avian embryos because early chick embryos fail to develop under hypoxic conditions.

To systematically evaluate how oxygen levels influence embryonic tissue development in the mouse, we performed droplet-based single-cell RNA sequencing (10x Genomics) on embryos cultured under normoxic and hypoxic conditions. Leveraging previously published, temporally highly resolved mouse single-cell whole embryo atlas data ^46^, we were able to identify the embryonic cell states expected during gastrulation, across both conditions. These included neuroectodermal derivatives (forebrain, midbrain, hindbrain, and spinal cord), mesodermal populations (paraxial, intermediate, and lateral plate mesoderm), endoderm, primordial germ cells, neural crest cells, and extra-embryonic tissues such as yolk sac (Figure S6A-B). The fidelity of tissue identity across both culture conditions highlights the capacity of our *ex-utero* system to support coordinated morphogenesis and lineage specification.

While the overall cellular composition remained broadly conserved between hypoxic and normoxic embryos (Figure S6B-C), differential gene expression analyses uncovered tissue-specific responses to oxygen availability (Figure S6D-F). Consistent with other contexts, elevation of oxygen induces significant downregulation of glycolysis pathway regulators, such as *Gapdh*, *Hk2*, *Slc2a1*, *Ldha* and *Aldoa* (Figure S6D). We further leveraged the full transcriptome of each cell to infer their stages, by aligning each cell to the reference atlas and observing the distribution of cell stages of its k-nearest neighbor references. The inferred stages of both conditions fall close to the expected 60-66 hour post-gastrulation (E8.5-E8.75) range (Figure S6F-G), with a slight overall delay in the relative developmental progression observed for embryos grown in normoxic conditions. Notably, some cell states exhibit more clear differences in developmental progression between hypoxic and normoxic conditions, with lateral plate mesoderm derivatives on average exhibiting potentially slight transcriptional acceleration (Figure S6G). This analysis, thus, highlights effects of oxygen levels on a variety of embryonic tissues. Most importantly in the current context, expression of canonical hindlimb specifiers such as *Pitx1*, *Tbx4*, together with Hox9 paralogues, were markedly upregulated in normoxic conditions, providing additional validation of the HCR results (Figure S6E; Figure 5). These findings substantiate the model that increased oxygen availability at later stages (as the placenta matures) selectively advances developmental timing in hindlimb field tissue.

We then asked whether the precocious expression of *Tbx4* in normoxia-cultured mouse embryos would lead to an acceleration of limb bud initiation. To that end, we examined the status of EMT in normoxia-cultured embryos after 48 hours, the equivalent of 66-hour post-gastrulation (E8.75). Using F-actin and LAMININ polarization as markers, we found that hindlimb embryos cultured in normoxia condition show a significantly higher proportion of mesenchymal cells in the hindlimb field (Figure 5H-I). Moreover, we found a correlation between *Tbx4* expression and EMT status, such that regions with higher *Tbx4* expression display more multi-layered mesenchymal cells (Figure S5D). Together these results suggest that higher oxygen levels accelerate hindlimb development, and support a model where maternal hypoxic conditions in the early developing mouse embryo act to specifically delay hindlimb development.

### Maternal hypoxia represses Tbx4 expression through *cRel* and *Hif1a*

We next sought to understand the downstream pathways mediating hypoxia’s repressive effect on *Tbx4* expression. While high oxygen culture normally induces a metabolic transition characterized by the downregulation of glycolysis and upregulation of oxidative phosphorylation (OXPHOS), in our context we found that neither the pharmacological inhibition of glycolysis nor the activation of OXPHOS was sufficient to induce the expression of Tbx4 (Figure S7A-D). These results ruled out the possibility that the shift in temporal expression of Tbx4 we observed is due to metabolic alterations.

To explore whether hypoxic conditions modulate the timing of *Tbx4* activation by regulating NFKB transcription, we performed qPCR on dissected hindlimb field tissues from embryos cultured under normoxia or hypoxia conditions. Consistent with the bulk RNA-sequencing analyses, among the NFKB family members, only the transcripts of *Rela*, *cRel* and *Nfkb1* can be readily amplified (Figure 5J-I, Figure S8A). Moreover, we found that in both qPCR and scRNA-seq analyses, the hindlimb tissues cultured in hypoxic conditions showed significant elevation of *cRel*, although non-significant expressional change of *Rela* and *Nfkb1* relative to the normoxia condition (Figure 5J, Figure S8A). These findings substantiate the hypothesis that transitioning from hypoxia to normoxia triggers the downregulation of *cRel*, thereby activating *Tbx4* expression.

To better understand how oxygen and *cRel* regulate *Tbx4* expression, we sought to dissect the regulatory relationships between *cRel* and the key transcriptional regulator of cellular responses to hypoxia, Hypoxia-inducible factor 1α (*Hif1a*). *Hif1a* is rapidly induced upon oxygen depletion to orchestrate adaptive gene expression programs. To determine whether oxygen-dependent regulation of *Tbx4* is mediated by *Hif1a*, we examined *Tbx4* expression in contexts where *Hif1a* levels were increased or decreased. We found that *Hif1a* overexpression in chicken hindlimb field induced significant downregulation of *Tbx4*, phenocopying the effect of *cRel* (Figure S9A-C; Figure 3D-H). Conversely, we generated *Hif1a* knockout embryos using the ES cell-embryo chimera approach, based on the design of a previously established *Hif1a* knockout mouse model^47^ (methods). Both ES cell colony genotyping and immunofluorescence in chimeric embryos confirmed the complete loss of HIF1A protein in all embryonic cells (Figure S9E-H). By E8.5, while wild-type embryos exhibited low *Tbx4* expression, *Hif1a* knockout embryos displayed a marked upregulation of *Tbx4* in the hindlimb field (Figure S9I-J). These findings indicate that *Hif1a* depletion phenocopies *cRel* loss-of-function, allowing premature *Tbx4* activation and positioning *Hif1a* as a key mediator of oxygen-dependent repression of *Tbx4*.

To further delineate the epistatic relationship between *cRel* and *Hif1a* in repressing *Tbx4*, we manipulated the expression of each factor individually and assessed the impact on the other. We observed that depletion of *Hif1a* did not significantly alter *cRel* expression. In contrast, modulation of cRel directly affected *Hif1a* levels: overexpression of *cRel* in the developing chicken hindlimb led to increased *Hif1a* expression, whereas *cRel* knockdown resulted in reduced *Hif1a* levels (Figure S10A–H). These findings indicate that, although both *cRel* and *Hif1a* are activated by hypoxia, *cRel* further promotes *Hif1a* expression, consistent with previous observations in other biological contexts^40^.

Overall, our results support a model where the maternal hypoxia to normoxia transition leads to downregulation of *cRel* and *Hif1a*, which allows the activation of *Tbx4*, and hence relieving a block on initiation of limb development in mouse embryos (Figure 6); and - from a broader perspective - explaining a critical aspect of heterochrony in limb formation between certain mammalian and avian species.

**Figure 6.**
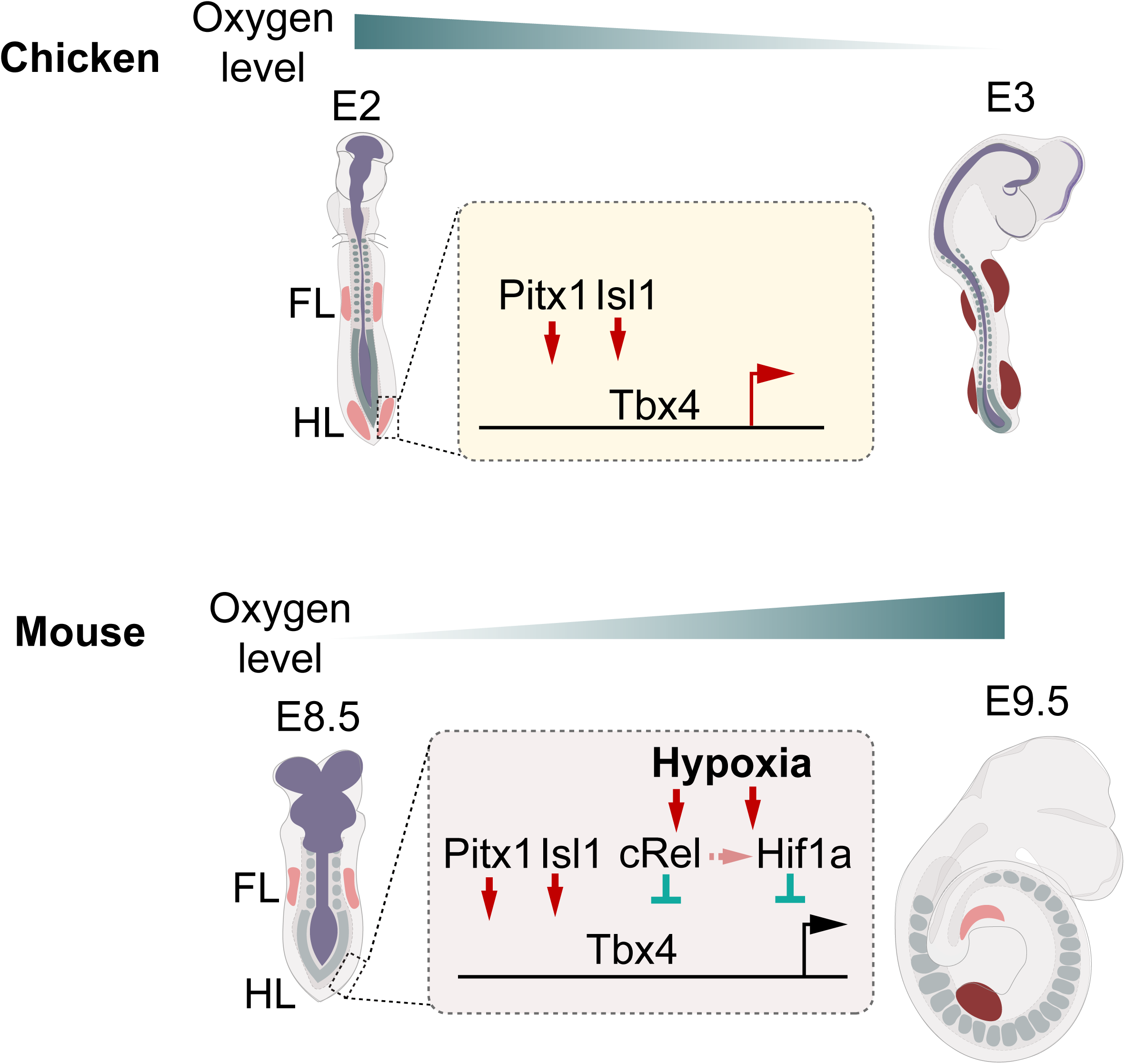
Summary model. Forelimb and hindlimb develop at different times from the limb initiation stage in the mouse embryo but much less so in the chicken, our results show that the limb initiation heterochrony in the mouse is mediated by the hypoxia environment that the mouse embryo exposes to in the limb field specification stage. The hypoxia environment inhibits limb initiation by delaying the hindlimb initiation factor such as Tbx4. Hypoxia’s effect on tbx4 expression is at least partially mediated by NFKB transcription factors such as cRel, and Hif1a. Altogether, our findings reveal the first example of the mechanisms behind limb initiation heterochrony and suggest a novel role of the maternal environment in directing the timing of embryonic development during post-implantation stages.

## Discussion

Despite growing interest in the study of developmental timing^43–45^, mechanistic understanding of heterochronic change remains limited. Our study suggests a model explaining the observed heterochronic delay (post-displacement) in the onset of hindlimb formation in mouse embryos relative to chicken by hypoxia (Figure 6). Avian species primarily obtain oxygen through a gas chamber and passive transport through the eggshell, which are limited in their oxygen-carrying capacity. As such, embryonic tissues gradually become hypoxic as cell numbers and tissue size increase ^41^. In contrast, mammals initially develop under hypoxic conditions due to the inaccessibility of embryonic tissues to oxygen. Subsequently, the maturation of placental functions allows efficient maternal-fetus oxygen exchange through allantois and blood circulation, and thus allows the transition to normoxia around mid-gestation stage. Mammalian embryos have evolved to exploit this increase in available oxygen as a cue to alter the timing of hindlimb formation relative to that seen in bird embryos. In particular, in the chick embryo *Pitx1* and *Isl1* activate *Tbx4* expression within 2-hour of their expression. But in the mouse embryo, *Tbx4* (but not its functional forelimb equivalent *Tbx5*) has been brought under the repressive influence of the NFKB family members *cRel* and *Rela*, which have been co-opted for this purpose by being up-regulated under hypoxic conditions. As a consequence, although expression of *Pitx1* and *Isl1* is activated in the hindlimb field at an equivalent developmental time as observed in the chick, they are unable to induce *Tbx4* expression for 18-22 hours, until oxygen levels rise and hence NFKB activity is diminished. Such differential temporal correlation between the expression of *Tbx5*, *Pitx1*, *Isl1* and *Tbx4* suggests that the difference in relative timing of limb bud initiation between mammals and birds derives from a hindlimb delay, rather than forelimb acceleration.

Our results show that that limb heterochrony in mammals (mouse) starts at the very first step of limb development, when *Tbx4* induces EMT. This contrasts with the delay in forelimb (wing) development in emu embryos, in which the reduced forelimb growth rate relates not to the limb initiation, but rather to changes in post-initiation cell proliferation ^33^. In this case, *Tbx5* expression occurs on time, but the expression of *Fgf10*, a key regulator of limb progenitor proliferation is reduced. This distinct mechanism is required in emus since their embryos, like chicks, lack an early hypoxic phase, and highlights the diverse levels of the limb developmental program that can be exploited by evolution to alter the tempo of limb outgrowth.

The delay in hindlimb development is even more pronounced in marsupial embryos, where it persists for longer period of time compared to placental mammals. As a result, the precocially born fetus displays an anatomically matured forelimb, capable of functioning in climbing to the pouch, but a still underdeveloped hindlimb^3^. Both forelimb acceleration and hindlimb delay have been posited as being responsible for limb heterochrony in marsupials^48–50^. Indeed, multiple mechanisms, involving a number of different levels of the limb morphogenesis program, may be involved in regulating this more extreme case of limb heterochrony. In this context, it is worth noting that, in spite of being much more extreme at later stages, the severity of marsupial limb heterochrony is similar to that of the mouse embryo at the limb initiation stage ^51^. Moreover, in the grey short-tailed opossum (*Monodelphis domestica*), both *Tbx5* in the forelimb and *Pitx1* in the hindlimb are detectable at least half a day earlier than *Tbx4* ^48^, as is the case in mice. These results at least raise the possibility that the mechanism we identify here as underlying eutherian hindlimb heterochrony may also be utilized in marsupial embryos as one of the adaptive changes contributing to their more severe hindlimb developmental delay. If so, then limb heterochrony in placental mammals might be an evolutionary vestige from the non-placental common ancestor of both marsupials and eutherian mammals, rather than having arisen *de novo* within the placental stem lineage.

One of the most intriguing findings of our study is the role of hypoxia in regulating the timing of hindlimb initiation. The transition from oviparity to viviparity dramatically alters the developmental environment from avian to mammalian species, with significant differences in oxygen levels as well as altering the level of accessibility of nutrients during the early embryonic development. As both parameters increase by placenta maturation, the level of oxygen can be viewed as a sensor of the level of nutrient availability. Mammalian embryos evolved to take advantage of this sensor to control the timing of development of less immediately required structures, such as the hindlimb, in an “energy trade-off strategy” to cope with limited nutrient availability.

When ectopically expressed, both REL and RELA blocked the expression of *Tbx4* and consequently the hindlimb bud formation. It is known that REL can competitively block the binding of RELA to the chromatin in other contexts ^52^. Therefore, although hypoxia does not significantly alter the expression of *Rela*, it is likely that *Rela* compensates for *cRel* activity only when *cRel* is depleted, explaining the weaker phenotype seen in *cRel* mutant embryos relative to that seen in embryos cultured in hypoxic conditions. In addition to *cRel*, we identified *Hif1a* as another hypoxia-responsive factor capable of mediating the repressive effect of hypoxia on Tbx4 expression. Our functional analyses indicate that *cRel* can promote *Hif1a* expression, at least to a modest extent, consistent with previous reports in other systems. Due to the technical limitations of combining *ex-utero* culture with transgenic mouse lines, such as *cRel* knockouts, our dataset lacks the statistical power to fully resolve the regulatory relationships among *cRel*, *Hif1a*, and *Tbx4*. Nevertheless, our results point to a complex interplay of hypoxia-responsive factors in regulating the timing of limb development.

Besides the activation of NFKB transcription factors described here, hypoxia can influence the activities of multiple pathways including protein synthesis rate, oxidative phosphorylation levels, and mitochondrial biosynthesis—factors known to regulate developmental timing in various contexts. Conversely, it is noteworthy that the NFKB pathway activities can also mediate cellular metabolism in other systems.

Overall, our studies provide the first investigation of the molecular mechanisms regulating the observed heterochrony in limb initiation, an important embryonic adaptation to *in-utero* mammalian development. The mechanism we elucidate represents a novel use of hypoxia as a trigger for temporal events in the embryo. In a broader sense, this provides new insight into how the maternal environment regulates developmental timing.

## Materials and Methods

### Animals and husbandry

Mice with various backgrounds were maintained and manipulated under the regulations of IACUC. Wildtype lines, including CD1, C57BL/6J and F1 (mice are purchased from Charles River. cRel mutant line was imported from Stephen Smale lab (UCLA) and was described before^53^. Tbx4 knockout line was purchased from Jackson Laboratory and was described previously^9^. Fertilized chicken eggs were ordered from Charles River farm and incubated at 38℃ with 40% humidity.

### Hybridization chain reaction (HCR)

Chicken embryos at HH8-HH23 stages and mouse embryos at E7.5-E10.0 stages were fixed in 4% PFA for 2h in room temperature or overnight at 4℃. Embryos were washed in PBST(PBS+0.1%Tween) for three times with 5mins each, and permeabilized in 70% ethanol in PBST for 2-5h at room temperature. The embryos were then washed by 37℃ pre-warmed washing buffer (Molecular Instruments co.) for 5mins and pre-hybridized in pre-warmed hybridization buffer (Molecular Instruments co.) for 30mins in 38℃ and then hybridized with hybridization buffer containing 60nM probes of targeted genes in 38℃ overnight. On the next day, the embryos were washed in pre-warmed washing buffer for two times with 30mins each time, followed by two times washes in 5xSSCT (5xSSC with 0.1% Tween) with 30mins each time. Embryos were pre-amplified in amplification buffer (Molecular Instruments co.) for 5mins at room temperature and then amplified in amplification buffer containing 48nM amplifiers matching the amplification sequences of each probe set. Before amplification, the amplifiers were heated in 95℃ for 90s and cooled down at room temperature for 30mins. The probes were designed following previous instructions.

### Chicken embryo electroporation and Early Chick (EC) culture

For HH13 lateral plate mesoderm electroporation, the chicken embryos were incubated for 50 hours, after which 5 ml albumin was removed from each egg and then windowed on the top. The electroporation was performed as previously described^54^. Briefly, plasmids with 1µg/ul mixed with FastGreen dye (0.2%) was injected into the coelom between two layers of lateral plate mesoderm of the forelimb or hindlimb and electroporated using BTX platinum needle L-shaped electrodes (45-0509). The electroporation was performed at 35 V, 5-ms pulse, 10-ms interval and 3-5 pulses. For HH4 chicken embryo electroporation, the eggs were cracked in 50-mm petri dishes and the albumin on the top of the embryo was gently removed using the blunt end of the forceps. The embryos were then transferred using pored filter papers (Whatman No.2) and was electroporated using Petri Dish Platinum Electrode using the following setting: 4 V, 50-ms pulse, 100-ms interval and 5 pulses. The embryos were then transferred to 35-mm petri dish coated with albumin+0.6% bacto agar supplemented with 15 mg glucose and 121 mM sodium chloride (NaCl). The embryos were then cultured in 38 ℃ for 24 hours.

### Immunostaining

The chicken or mouse embryos were first fixed in 4% PFA for 2h in room temperature or overnight at 4℃. The embryos were then dehydrated in 30% sucrose and embedded in 50% OCT in 15% sucrose solution and then processed with cryostat sectioning. The OCT in sections were washed by PBS for two times and then blocked by TNB solution (150 mM NaCl, 100 mM Tris-HCl (pH=7.5), 0.3% Triton X-100, 0.5% blocking reagent (Roche, 11096176001)) for 30mins at room temperature, and then incubated with TNB solutions containing primary antibodies in 4C for overnight. For Hif1a staining, primary antibody was diluted in PBS. The slides were then washed for 3×5mins in PBS and incubated with 2^nd^ antibodies for 1h at room temperature. After 2^nd^ antibody incubation, PBS or PBS containing DAPI and F-actin (diluted in 1:1000) were incubated for 5mins, followed by PBS wash for 5mins and mounted with Invitrogen™ Fluoromount-G™ Mounting Medium covered by cover slides. Dilutions for primary antibodies: Rabbit Anti-Laminin polyclonal antibody (Sigma-Aldrich, L9393), 1:500. Rabbit anti-HIF1a Polyclonal Antibody (Proteintech, 20960-1-AP), 1:100. Rabbit anti-phospho-Smad1 (Ser463/465)/ Smad5 (Ser463/465)/ Smad9 (Ser465/467) (D5B10) monoclonal antibody (Cell Signaling Technology, 13820S), 1:200. Secondary antibodies are diluted in TNB at 1:500 dilutions. List of antibodies include: donkey anti-Rabbit 488 (Jackson ImmunoResaerch, 711-545-152), donkey anti-Rabbit 568(Invitrogen, A10042), donkey anti-Rabbit 647 (Jackson ImmunoResaerch, 711-605-152).

### Plasmids construction

Enhancer:reporter constructs: for constructing enhancer:reporter plasmids, pTK-RetE1-Citrine construct (a gift from Tatjana Sauka-Spengler (Addgene plasmid # 127779)) was double digested by KpnII (NEB, #R3142) and XmaI (NEB, R0180S) enzymes to create pTK-Citrine. Enhancer candidate regions were PCR amplified from chicken genomic DNA, with 5’ and 3’ primers carrying 18-22nt homologous sequences overlapping with the upstream and downstream of KpnI and XmaI sites of pTK--Citrine construct. The enhancer candidate regions were then cloned into pTK-Citrine vector via Gibson assembly. Positive clones were selected by both enzyme digestion and DNA sanger sequencing. The plasmids were maxi-preped before electroporation. Primers used for cloning are listed in Table S2.

Overexpression construct: pCAGGS-T2A-IRES-ZsGreen construct was double digested with XhoI (NEB, R1046) and EcoRI (NEB, R3101) enzymes. The protein coding sequence of cREL were amplified from chicken HH23 forelimb and hindlimb cDNA. 5’ and 3’ primers carrying 18-22nt homologous sequences overlapping with the upstream and downstream of XhoI and EcoRI sites of pCAGGS-T2A-IRES-ZsGreen construct. The coding sequences of genes of interest were then cloned into the construct via Gibson assembly. Positive clones were selected by both enzyme digestion and DNA sanger sequencing. The plasmids were maxi-preped before electroporation. The following primers are used:

### *Ex-utero* culture and pharmacological treatments

E6.5 embryos were recovered from CD1 females mated with either CD1 or CB6F1 males, and were cultured in the medium containing 50% heat-inactivated rat serum (Biomed), 5% Fetal Bovine Serum 1× glutamax (GIBCO, 35050061), 100 units/ml penicillin and 100 μg/ml streptomycin (Biological Industries; 030311B) and 11mM HEPES (GIBCO, 15630056). Up to 3 embryos were placed in 12-well plate containing 2 ml medium each well. For each experiment, the embryos from the same batch were split into two groups and cultured for up to 2 days in 5% or 20% O_2_ levels. All pharmacological inhibitors or activators were added directly into the ex-utero culture medium. 2-DG was treated at 0.1mM, Dichloroacetate (DCA) was treated at 0.15%.

For JC-1 staining, the hindlimb field from ex-utero cultured embryos were dissected manually and disassociated as single cells and run through flow cytometry. The Grenn (FL-2) to Red (FL-3) ratios were counted and compared.

### Real-time quantitative PCR (RT-qPCR)

RNA was extracted from dissected hindlimb field of ex-utero cultured embryos from E6.5+1.5 days, under 5% or 20% O_2_, using the Arcturus PicoPure RNA isolation kit (Arcturus Bioscience). RT-PCR was performed using a StepOne Plus Real-time PCR machine (BioRad). The expression level was calculated using ddCT methods, normalized to a Atp5f2 PCR reaction and the endogenous control group. The primers used for RT-PCR are as below (all targeting mouse RNA sequences): Atp5f1-F: AGTTCCTTTACCCTAAGACTGGT; Atp5f1-R: TTCATGCTCGACTGCTTTACTT. cRel-F: ACAACAACCGGACATACCCG; cRel-R: GGTCTGCGTTCTGGTCCAA. RelA-F: AGCGCGGGGACTATGACTT; RelA-R: GCCCGGTTATCAAAAATCGGAT. Nfkb1-F: ATGGCAGACGATGATCCCTAC; Nfkb1-R: CGGAATCGAAATCCCCTCTGTT. Nfkb2-F: TGGCATCCCCGAATATGATGA; Nfkb2-R: TGACAGTAGGATAGGTCTTCCG. RelB-F: GTTCCAGTGACCTCTCTTCCC; RelB-R: CCAAAGCCGTTCTCCTTAATGTA.

### Making transgenic embryonic stem cells (ES)

Wild type E14 embryonic stem cells were transfected with two constrctus, that one codes super-piggybac transposases (NovoPro, V012800), and one contains piggybac LTRs enclosing three cassette: a) enhancer:reporter cassette that contains enhancer:Hsv-tk:Citrine; b) CMV:H2B-mCherry; 3) Eif1a: puroR. Two days after transfection, the positive clones were first selected by puromycin treatment at 1ug/ml. After a week, the cells were double selected by manually picking the single cell formed colonies that do not show obvious citrine signal and detectable mCherry signal. Multiple clones were mixed and cultured together.

### Making ES-embryo chimera

For making the chimera, F1 females (CB6F1, Charles River) were superovulated by PMSG injection on Day 1 and HCG injection on Day 3, and mated with F1 males to produce the embryos. The embryos were recovered at E1.5 and cultured until E2.5 in KSOM medium, on which day the zona pelluccida was removed by acid tyrode’s solution. The transgenic ES cells were trypsinized into single cells, and clamps with about 20 cells were aggregated with E2.5 mouse embryos before compaction. The chimeric embryos were then cultured in KSOM until E3.5, after which the chimeric embryos were transferred into the uterus of pseudo-pregnant CD1 females who were mated with sterilized CD1 males three days before the experiment. 20 chimeric embryos were transferred into each pseudo-pregnant female, and each experiment averagely 4 females were used for transfer.

### Generation of Hif1a knockout ES cells

The region of depletion for *Hif1a* locus (containing a start coding site of Hif1a) was designed aas a previous study^47^ (to keep the consistency, chromosome 12: 73973312 to 73973522). Four gRNAs targeting sequences upstream and downstream of the deletion region were designed using the CHOPCHOP website and transfected, together with Cas9 protein, using Lipofectamine CRISPRMAX Cas9 Transfection Reagent (Life Technologies Corporation, CMAX00001). Transfected ES cells were cultured for 48 h and then dissociated into single cells. Individual cells were isolated by mouth pipetting into 96-well plates for clonal expansion.

Once colonies reached approximately 70% confluency, genomic DNA was extracted for genotyping using primers flanking the deleted region: forward, 5′-ctttgttcatgttccatacaagtcc-3′; reverse, 5′-ccttcataagagcatattaatatgtgcac-3′. Homozygous deletions yielded a single band of ∼336–364 bp, whereas wild-type and heterozygous clones produced an additional band at 502 bp. Homozygous knockout ES cell clones were expanded in 6-well plates prior to aggregation with CD1 host 8-cell stage embryos for chimera generation. Deletion of Hif1a was further confirmed by immunostaining for Hif1a protein. gRNA sequences are as below: gRNA1: CTAATCTTTATACAGTAAACAGG; gRNA2: TAATCTTTATACAGTAAACAGGG; gRNA3: CTAGAGATGCAGCAAGATCTCGG; gRNA4: TCCTATATAATAATTTTACATGG.

### Bulk RNA-sequencing and analyses

Tissues corresponding to the area of forelimb and hindlimb of chicken embryos at HH12, and mouse embryo at E8.5 and E9.5 were finely dissected and disassociated as single cell in TrypLE buffer. The RNA from each tissue was extracted using RNeasy Kit (Qiagen). 1ng of RNA was used for smart-seq v4 RNA sequencing. RNA was sequenced in pair-end. For the analyses, RNA sequencing reads were mapped to chicken genome Galgal6, and mouse genome mm10 by hisat2. FPKM (Fragments Per Kilobase per Million mapped reads) of genes were calculated by featureCounts, differentially expressed genes were calculated by DESeq2.

### ATAC-seq identification of candidate CRE loci

The raw data deposited on GEO (GSE136775), as reported on Young et al (*Curr Biol* 2019 Nov 4;29(21):3681-3691.e5), was downloaded. After cleaning up and QC’ing the FASTQ files with TrimGalore (Krueger, 2015) and aligned them to the galGal6a (NCBI) genome using BWA (Li & Durbin, 2009), peaks were called with MACS2 (Zhang et al, 2008) using parameters recommended for ATAC-seq and genome coverage was determined using DeepTools (Ramirez et al, 2016). Differential accessibility between hindlimb, forelimb, and flank were calculated using the BDGDIFF function within MACS2 (signal cutoff: 10^2^).

### Single cell RNA sequencing & analysis

As described above, ex-utero cultured embryos in hypoxia (2-5% O2) or normoxia (21% O2) for two days were dissociated in TrypLE at 37 degrees for 10 minutes with pipetting every 5 minutes. The dissociated cells were mixed with recovery medium (10% FBS in low glucose DMEM) and spun down for 5 minutes at 40 rcf, followed by one time wash in PBS, and ultimately placed in PBS with DRAQ5 (ThermoScientific, Cat # 62251, 10 uM) and DAPI (1 ug/mL), and placed on ice. The samples were subject to On-chip Sort microfluidic chip sorter (On-chip Biotechnologies) to enrich for viable cells (DRAQ5+DAPI-). During the sort, it was ensured that the cells were in ice cold condition. The resulting sorted suspensions in 1X PBS and 1% BSA were put into separate lanes for GEM generation according to the 10X Chromium GEM-X Single Cell (Dual Index) protocol. To minimize further any sources of batch effect and have an internal control for genotype demultiplexing, an additional lane was run with the hypoxic and normoxic condition samples combined. Subsequent post GEM-RT cleanup, cDNA amplification, and 3’ gene expression library construction steps were performed in accordance with the manufacturer’s instructions. Sample indices were chosen from the 10X Dual Index Kit TT Set A (PN-1000215). Sequencing was performed on a NovaSeq X flowcell at the Biopolymers Facility at Harvard Medical School according to the recommended 10X sequencing specification.

Generation of the count matrix (cellranger v9.0.0), SNP detection (cellsnp-lite v1.2.3), and genotype-based demultiplexing (vireoSNP v0.5.8) were all performed on the Harvard Medical School O2 computing cluster. To facilitate the computational time involved in genotype inference, all cells from the three libraries were randomly partitioned into four sets of ∼40,000 cells that overlap 50% overlap with another partition, and vireo was run with 40 randomly generated seeds to find the best log-likelihood fit. The number of expected genotypes were set to 24 (N=24) to obtain inferred genotypes. Given all cells were at least two times inferred from separate partitions, the genotype for each partitions could be stitched back for the whole dataset. Additional filtering was conducted to identify the cellular barcodes that conform to the expected pattern of exclusion for a specific library (presence exclusively in hypoxic condition library or normoxic condition library, with common presence in the shared library). These final genotype inferences were used to reconstruct 9 embryos from hypoxic conditions and 10 embryos for normoxic conditions. Ambiguous cellular barcodes (genotype inferences that were found in all three libraries, potential doublets that “cross-over” between genotypes in two overlapping partitions) were discarded. After generation of Seurat objects, downstream analysis using the Seurat package was performed in R^55^ (Hao et al., 2023; Hao et al., 2021; Stuart et al., 2019; Butler et al., 2018; Satija et al., 2015).

The embryonic gastrulation atlas (Imaz-Rosshandler et al. 2024) has been found to have some mis-annotations of the mesenchymal lineage. As such, the data was subsampled to contain E7.75 to E9.25 samples and were re-processed for batch-correction (harmony reference) and graph-based clustering using the Leiden algorithm (resolution=0.8) that retains most of the mapping from the original annotation but fixed the paraxial, lateral plate mesoderm lineage and incorporated proper separation of the limb fields into forelimb and hindlimb clusters. The resulting reference annotation was used for LabelTransfer procedure in Seurat (v5) to label cells of the ex-utero dataset. The number of inferred cell state annotations were normalized against the number of deconvoluted genotype (embryo), and this relative fraction was compared between the two conditions. The predicted stage was derived from a subsampled reference dataset where all the time points from E7.75 to E9.25 have equal numbers (by random subsampling) and the stage values were changed to numerical value, and with the anchored label transfer procedure, the weighted k-nearest mean value of stages was associated with the query cells. Gene expression visualization was conducted by pseudobulking by embryo or embryo & predicted cell state. T-test with Benjamini Hochberg multiple testing adjustment was performed for differential gene expression. The size of the pseudobulk was only considered for filtering too small pseudobulk sets, given the average level of the gene in question (high expressing genes for Figure S6D, >100 cells; mid-expressing genes for S6E, >400 cells; for low-expressing genes for S8A, >2000 cells). The stage distribution analysis, the default parameters for ‘geom_density‘ function in ‘ggplot2‘ package was used to display individual embryo and whole condition distributions. Kolmogorov-Smirnov test was performed on the stage distributions between hypoxic and normoxic conditions for difference.

## Material availability

All materials are available through reasonable requests to the corresponding author.

## Acknowledgment

We thank all members in Cepko-Tabin labs and Susan Dymecki lab, and Dr. Emma Farley, Dr. Scott Edwards for giving constructive comments on the project. We thank Dr. Macolm Logan and Dr. Ben Steventon for helpful discussions at the initial phase of the project. We also thank Dr. Nandan Nerukar and Dr. Marianne Bronner for the training of early chick culture experiments, Dr. Lina Du from mouse engineering core at Dana-farber with the help of embryo transfer, and Micron imaging core especially Dr. Paula Montero Llopis for imaging assistance.

## Author contribution

Conceptualization: M.Z; Formal analysis: M.Z, R.C.P. Funding acquisition: C.J.T., M.Z; Investigation: M.Z, C.L, C.J.T; Methodology: M.Z, C.J.T; Visualization: M.Z; Supervision: C.J.T.; Writing-original draft: M.Z, C.J.T.; Writing-review & editing: M.Z, R.C.P, C.L, C. J.T.

## Data and code availability

The bulk RNA-sequencing data, as well as thd single cell RNA-sequencing data has been deposited to NCBI GEO database (GSE298623 for bulk RNA seq; GSE298624 for single cell RNA-seq, in embargo until next year, can be released prior to publication upon requests). Original images for figures are deposited to figshare (https://figshare.com/account/home#/projects/251516). All other raw data are available upon reasonable request to the corresponding author. All processing code for the scRNA-seq analysis are available at https://github.com/chlee-tabin/limb-heterochrony.

---

**Figure S1.**
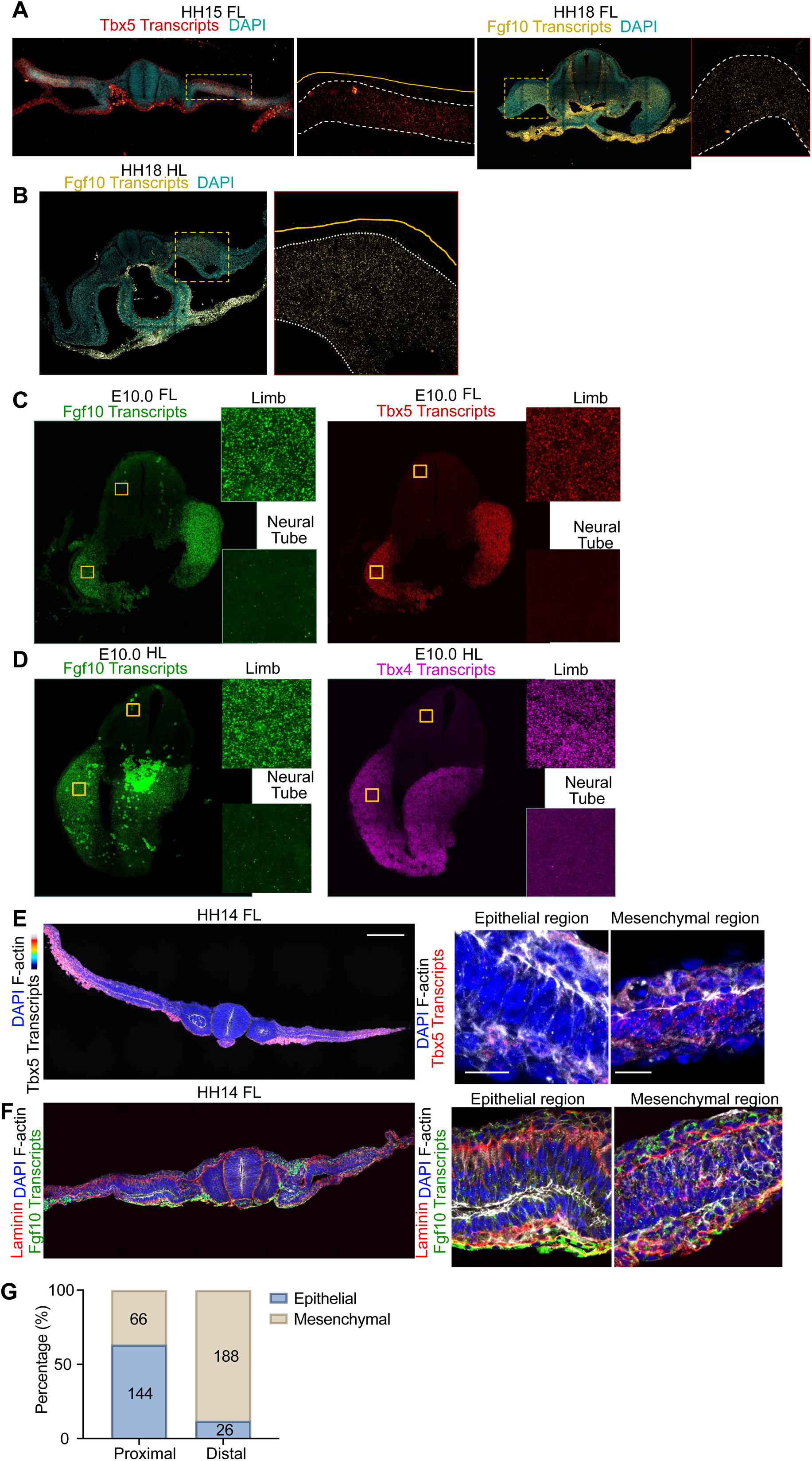
Expression of Tbx5 and Tbx4, but not Fgf10, correlate with the timing of EMT. (**a**-**b**) Visualization of Tbx5 in HH15 chicken forelimb, and visualization of FGF10 expression by HCR in forelimb (**A**) tissues and hindlimb (**B**) tissues of the chicken embryos at HH18. White dotted lines indicate magnified regions where yellow dotted lines indicate the region of LPM. N=5 embryos, N=4 experiments. (**C**) Visualization of Fgf10 and Tbx4 expression by HCR in E10.0 mouse forelimb tissue. (**D**) Visualization of Fgf10 and Tbx4 expression by HCR in E10.0 mouse hindlimb tissue. For **C**-**D**, yellow squares indicate magnified regions. N=6 embryos. N=3 experiments. (**E**) Correlation between TBX5 expression in chicken HH14 forelimb tissues by the Tbx5 transcripts visualized by HCR and F-actin and DAPI staining. Higher Tbx5 expression correlates with higher proportion of mesenchymal state. Quantification shown in Fig.1H. (**F**) No correlation between FGF10 transcripts levels and EMT in chicken HH14 forelimb tissues, as shown by the level of Fgf10 transcripts (visualized by HCR) and Laminin, F-actin and DAPI staining. For **E**-**F**, yellow dotted lines indicate proximal region where cells are mostly epithelial, dotted lines indicate distal region where cells are mostly mesenchymal. N=2 experiments, N=4 embryos. (**G**) Quantification of epithelial and mesenchymal cells in the medial versus lateral regions of HH14 (forelimb) and HH15 (hindlimb) chicken embryos. Regional divisions are defined by the midpoint of the lateral plate mesoderm. Data are presented as bar graphs showing mean ± S.E.M. N=3 embryos from N=3 experiments. Scale bars for overview images, 150μm; scale bars for magnified images, 15μm.

**Figure S2.**
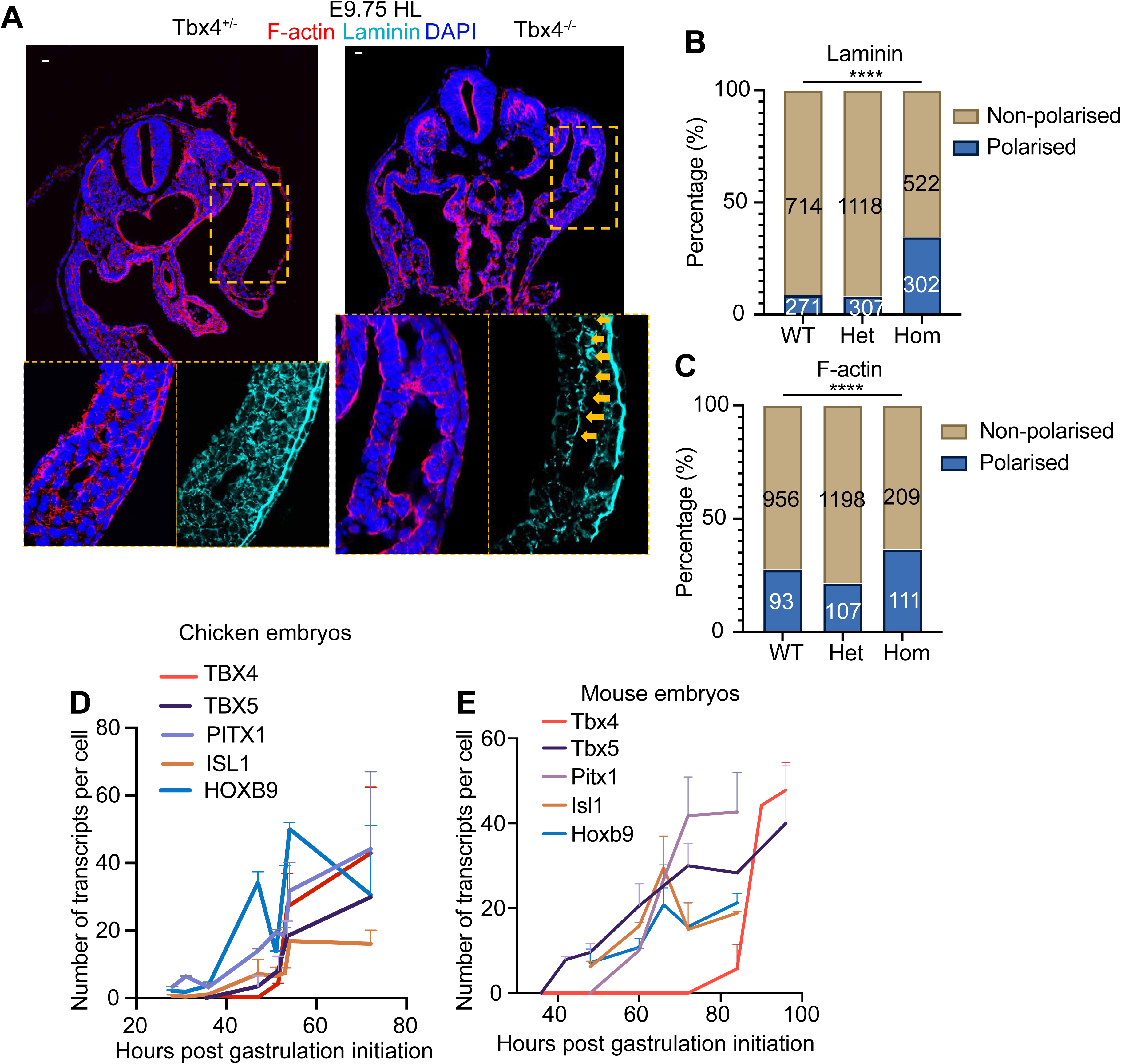
Tbx4 expression regulates hindlimb EMT and the delay of its expression is not regulated by the timing of Hoxb9, Pitx1 and Isl1 expression. (**A**) Cellular states in the hindlimb tissues of Tbx4 heterozygous and homozygous knockout embryos at E9.75. Dotted lines indicate magnified regions and the yellow arrows indicate epithelial cells that fail to undergo EMT on time and retained in the basal of hindlimb tissues in Tbx4 homozygous knockout embryos. N=6 embryos for Tbx4 heterozygous mutants and N=3 embryos for Tbx4 homozygous mutants. (**B**-**C**) Quantification of cells showing laminin (**B**) or F-actin (**C**) polarization as indications of epithelial state. The homozygous knockout embryos show higher proportion of epithelial cells comparing to wildtype (WT) or heterozygous embryos (Het). ****p<0.0001, Fisher’s exact test. Data shown as bar charts with numbers in each bar represents the number of cells examined. N=3 embryos for WT, N=6 embryos for Het, and N=3 embryos for Hom. N=2 experiments. (**D**-**E**) Quantification of transcripts levels of Pitx1, Isl1 and Hoxb9 across HH8-HH18 in chicken embryos (**D**) and E7.5 to E10.0 in mouse embryos (**E**). The expression curves are aligned to that of Tbx5 and Tbx4, which reveals that while Hoxb9/Pitx1/Isl1 expression is almost concomitant with Tbx5 and Tbx4 in the chicken embryo, in the mouse embryo they only expression together with Tbx5, and Tbx4 expression is delayed. Data is shown as mean ± S.E.M. Scale bars for overview images, 150μm; scale bars for magnified images, 15μm.

**Figure S3.**
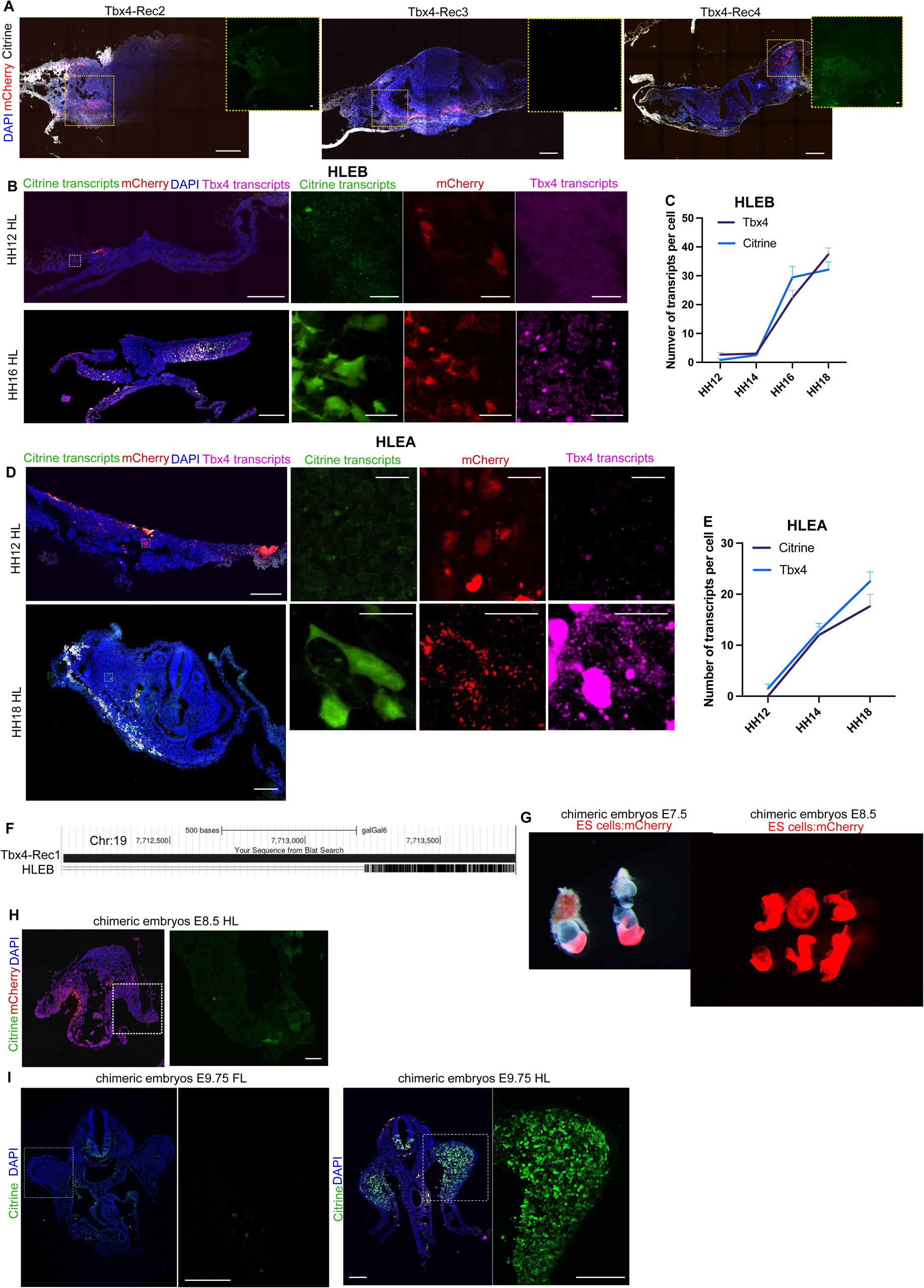
Chicken Tbx4 enhancer is conserved to that of mouse Tbx4 enhancer HLEB. (**A**) Enhancer:reporter constructs carrying Tbx4 enhancer candidate, Tbx4-Rec2, Tbx4-Rec3 and Tbx4-Rec4 were electroporated into the HH13 chicken embryos and the enhancer activity was examined at HH18 by Citrine signal. mCherry and DAPI were also revealed as electroporation and cell position control. Besides Tbx4-Rec1 (shown in Fig.3D), none of the other Tbx4 enhancer candidates show activities. Dotted lines indicate magnified regions. N=6 embryos for each construct, N=2 experiments. (**B**) Representative images of chicken hindlimb tissues electroporated with the HLEB:Citrine enhancer-reporter construct, either co-electroporated with pCMV-mCherry (for embryos at HH12–14) or without (for embryos at HH16–18). Sections were collected from stages HH12 to HH18 and stained with two marker combinations: embryos at HH12–14 were stained with Citrine HCR, mCherry, and TBX4 HCR; embryos at HH16–18 were stained with Citrine, Citrine HCR, and TBX4 HCR. Quantifications are presented in panel **C**. (**C**) Quantifications of Citrine transcripts number, reading out HLEB activity, against TBX4 transcripts number from cells in hindlimb tissues of chicken embryos shown in **B**. (**D**) Representative images of chicken hindlimb tissues electroporated with the HLEA:Citrine enhancer-reporter construct, either co-electroporated with pCMV:mCherry (for embryos at HH12–14) or without (for embryos at HH16–18). Sections were collected from stages HH12 to HH18 and stained with two marker combinations: embryos at HH12–14 were stained with Citrine HCR, mCherry, and TBX4 HCR; embryos at HH16–18 were stained with Citrine, Citrine HCR, and TBX4 HCR. Quantifications are presented in panel **E**. (**E**) Quantifications of Citrine transcripts number reading out HLEA activity, against TBX4 transcripts number from cells in hindlimb tissues of chicken embryos shown in **B**. (**F**) Alignment of sequences between Tbx4-Rec1 and HLEB. 3’ sequence of Tbx4-Rec1 show conservation to HLEB. The alignment is visualized through UCSC genome browser. Conservation score, 312. (**G**) Stereoscope images ES-embryo chimera dissected at E7.5 and E8.5. The transgenic ES cells are labelled with mCherry signal. Full penetration of the donor ES cells is achieved. Scale bars for all images, 150μm.

**Figure S4.**
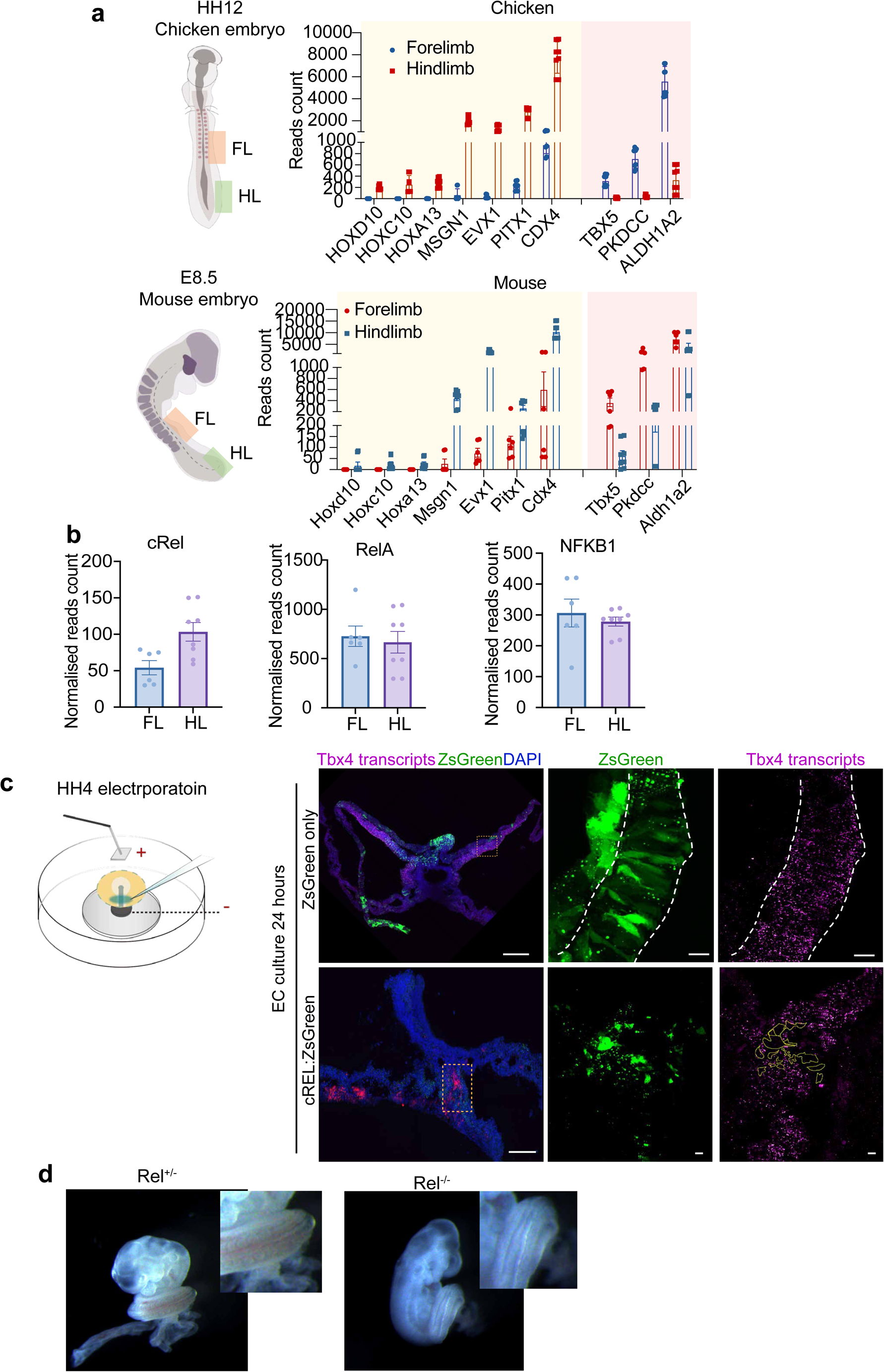
cRel represses the expression of Tbx4. (**A**) Bulk-RNA sequencing faithfully captures the transcriptional signatures of forelimb and hindlimb tissues at HH12 chicken embryo and E8.5 mouse embryo. (**B**) Expression levels of *cRel*, *RelA* and *Nfkb1* in E8.5 mouse embryos and in forelimb and hindlimb fields from bulk-RNA seq shown in **A**. (**C**) Electroporation of CAG:cRel:ZsGreen overexpression constructs into the mesoderm and ectoderm of gastrulating chicken embryos at HH4 led to significant downregulation of Tbx4 in hindlimb field. Construct containing only ZsGreen was electroporated as a control. (**D**) *cRel* knockout embryos do not accelerate hindlimb bud formation at E9.5 and show similar hindlimb thickness as heterozygous littermates. N=4 embryos for heterozygous and N=4 embryos for homozygous. N=2 experiments. (**E**) Quantification of the ratio of hindlimb/forelimb size in heterozygous and homozygous *cRel* mutants at E9.5. Data shown as mean± S.E.M. Each dot indicates an embryo. Scale bars for overview images, 150μm; scale bars for magnified images, 15μm.

**Figure S5.**
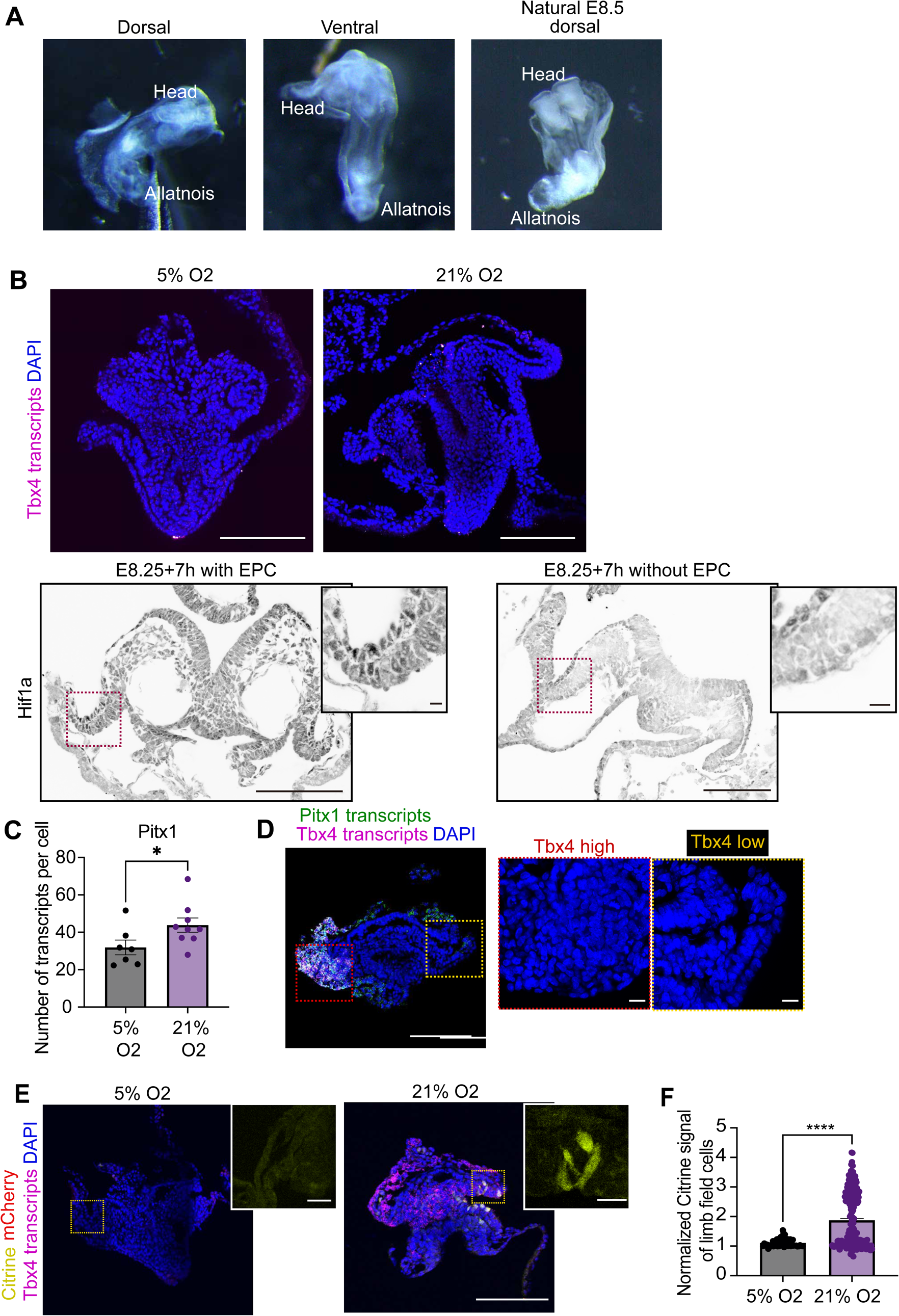
*Ex-utero* culture of embryos with prematurely elevated oxygen level show accelerated expression of hindlimb initiation regulators. (**A**) Gross morphology of embryos cultured in 5% O_2_ using *ex-utero* culture method shown in Fig.5**d** and natural E8.5 mouse embryo. The morphologies of ex-utero cultured and natural E8.5 mouse embryos are similar expect that there were occasional defects in somite morphogenesis. N=22 experiments. (**B**) Culturing embryos from E7.5 to E8.5 does not induce expression of Tbx4, and removal of EPC facilitates higher efficiency of oxygen transportation as indicated by the downregulation of Hif1a in the posterior embryonic tissues of E8.25 embryos cultured ex-utero for 7h. Red dotted lines indicate magnified regions. N=2 sections from N=2 embryos. N=2 experiments. (**C**) *Ex-utero* culture of embryos (protocol shown in Fig.5D) in normoxia led to significant upregulation of Pitx1 in hindlimb tissues. *p<0.05, Student’s t test. Each dots indicate one embryo. N=8 experiments for both 5% and 20% O_2_. (**D**) Chimeric embryos carrying Tbx4-Rec1:Citrine transgenic ES cells were ex-utero cultured from E6.5 over the course of two days in hypoxia or normoxia, and stained with Citrine, mCherry, Tbx4 transcripts and DAPI. Magnified regions indicating the hindlimb field cells. (**E**) Quantifications of Citrine signal in cells from embryos shown in **D**. each dots represents a quantified cell. N=2 experiments, N=4 embryos for both hypoxia and normoxia conditions. (**F**) Expression of Tbx4 correlate with cell number and epithelial/mesenchymal state as revealed by the expression of *Pitx1* and *Tbx4* transcripts (by HCR) and the staining of DAPI. Red dotted line indicates region that highly express Tbx4 (more cell numbers and multi-layer of cells) and yellow dotted line indicates the region that show no Tbx4 expression (less cell numbers and mono-layer of cells). Scale bars for overview images, 150μm; scale bars for magnified images, 15μm.

**Figure S6.**
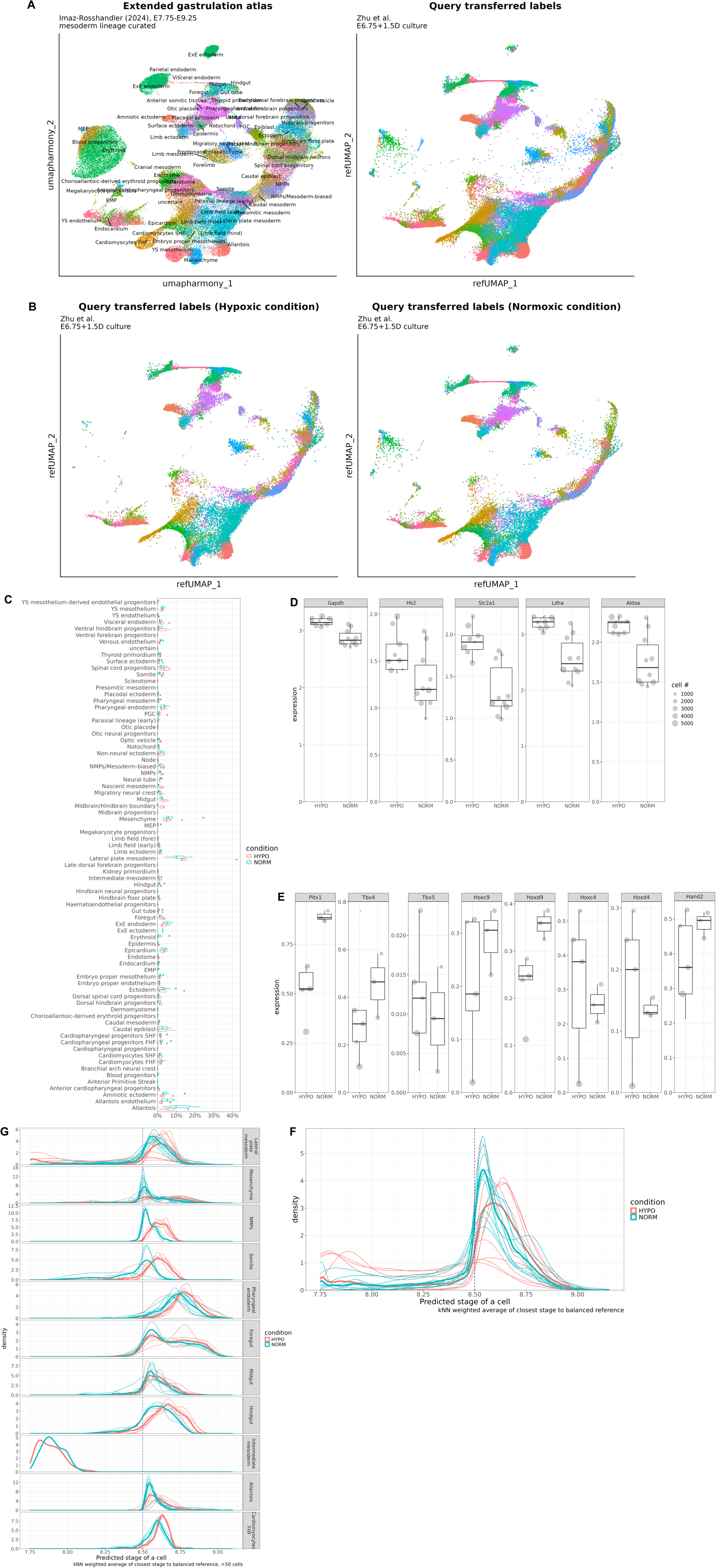
Single-cell RNA sequencing reveals tissue-specific transcriptional responses to hypoxia. (**A**) UMAP plot of re-clustered extended gastrulation atlas by Imaz-Rosshandler (2024), for stages from E7.75 to E9.25 (Extended gastrulation atlas), and the ex utero cultured mouse embryos from this study in the same UMAP embedding (Query transferred labels). The colors of the right panel matches with the left panel (**B**) Same UMAP embedding split by the hypoxic and normoxic conditions. (**C**) Cell state composition analysis. Fractions of all curated cell state annotations against the total number of cells from the same embryo, boxplots colored by hypoxic and normoxic conditions (9 and 10 embryos respectively). (**D**) Known target gene expression of Hif1a in pseudobulk, split by hypoxic/normoxic conditions. Hypoxic=9 embryos, Normoxic=10 embryos. Dot size represent the number of cells to constitute the pseudobulk. All genes have stistically significant differential expression (p-value < 0.05 by Benjamini-Hochberg multiple testing adjustment) (**E**) Hindlimb markers, forelimb markers, and lineage progression related marker gene expression in the lateral plate mesoderm annotated cells. Hypoxic=4 embryos, Normoxic=3 embryos that had sufficient number of lateral plate mesoderm cell type. Dot size represent the number of cells that constitute the pseudobulk measurement. Despite smaller sample size and coverage, Pitx1, Hoxd9 show statistically significant differential expression (p-value < 0.05 by Benjamini-Hochberg multiple testing adjustment) (**F**) Inferred stage distribution of individual embryos (fine curve) and aggregated condition (thick curve) (**G**) Cell state specific inferred stage distribution between hypoxic and normoxic conditions with thick curve showing aggregate distribution, and thin curve showing individual embryos. Given the extremely low cell depth of the whole embryo partitioned into individual cell types and embryo, most of the Kolmogorov-Smirnov test was found not significant except Hindgut (p=0.012), NMPs (p=0.026), Cardiomyocytes (First heart field) (p=0.039), Mesenchyme (p=0.044) and Somite (p=0.045), with Lateral plate mesoderm showing acceleration (difference in mean stage between hypoxic and normoxic condition of 0.082, p=0.054).

**Figure S7.**
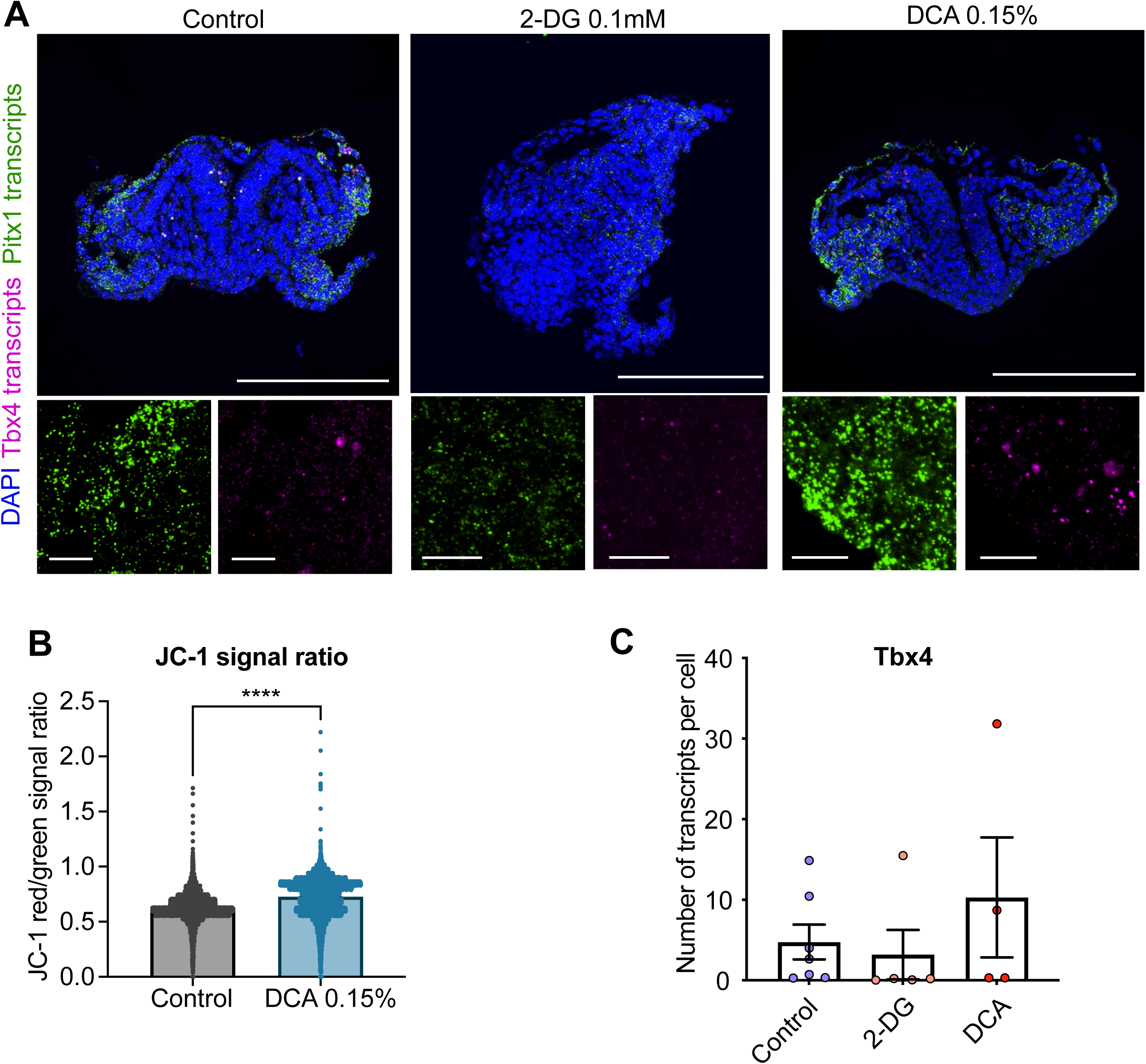
Alterations of glycolysis and oxidative phosphorylation activities corresponding to changes between hypoxia and normoxia did not change the timing of Tbx4 expression. (**A**) Representative images of embryos cultured ex-utero in hypoxia, and treated with water (control), glycolysis inhibitor (0.1mM 2-DG), and oxidative phosphorylation activator (0.1% DCA) for 48 hours from E6.5, and stained with Pitx1 HCR, Tbx4 HCR and DAPI. Magnified images represents the hindlimb field tissues. (**B**) Quantifications of mitochondria membrane potential as a readout of oxidative phosphorylation revealed by JC-1 signal ratio (Red/Green). Data generated by flow cytometry of single cells from dissected hindlimb tissues. Data shown as mean SEM, each dot represents one cell, N=2 experiments. (**C**) Quantifications of number of Tbx4 transcripts in hindlimb field cells from ex-utero cultured embryos treated in conditions shown in **A**. Each dot represents one embryo. N=3 experiments. ns, not statistically significant. One way ANOVA. Scale bars for overview images, 150μm; scale bars for magnified images, 15μm.

**Figure S8.**
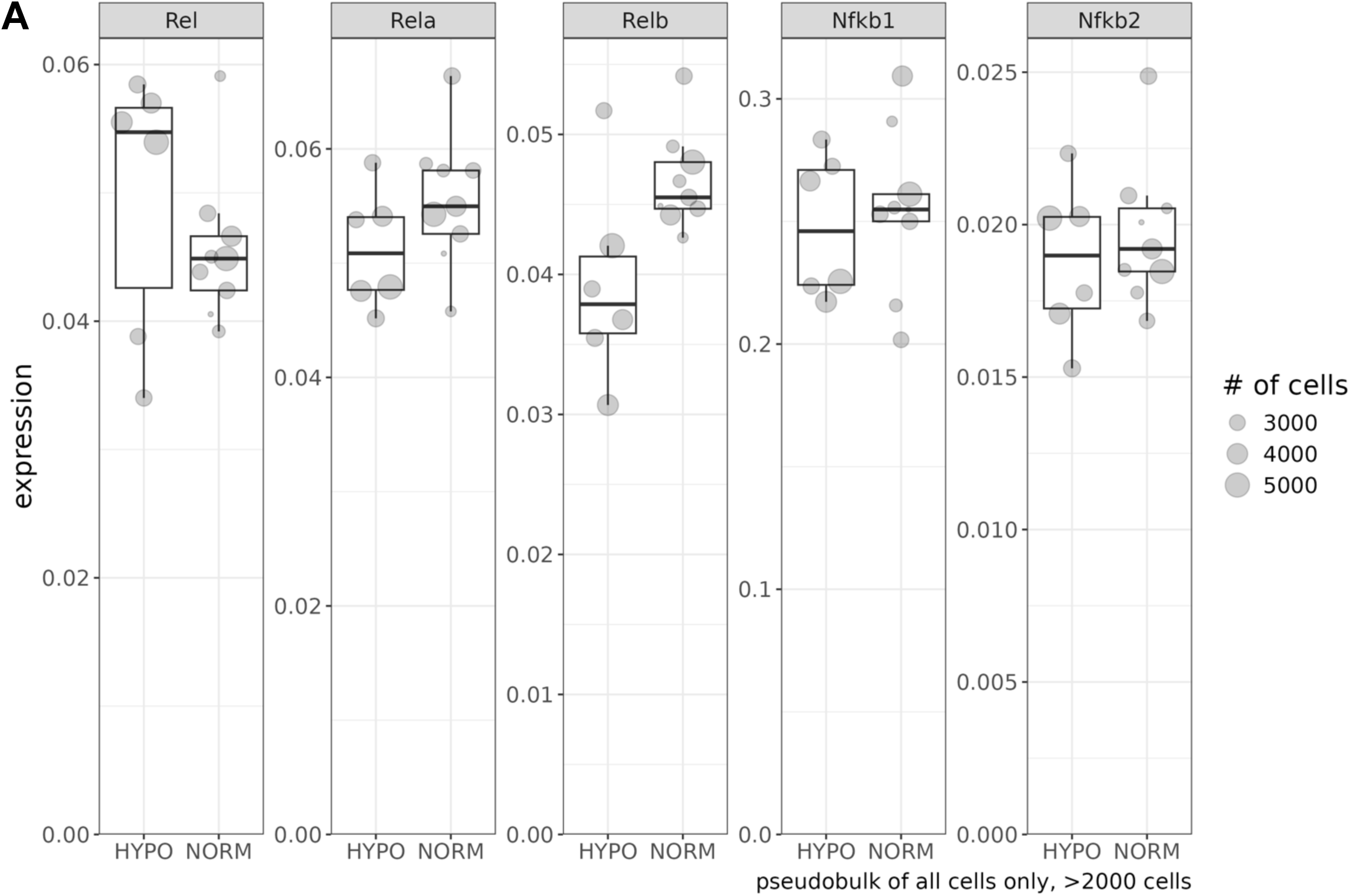
Hypoxia inhibits Tbx4’s expression partially through cRel. (**A**) cRel related paralog expression level in hypoxic and normoxic conditions from the single cell RNA-seq analysis. Due to their more general low level of expression more stringent filters were employed (the pseudobulk sample should have more than 2,000 cells). Hypoxic=6 embryos, Normoxic=9 embryos. Dot size represents the number of cells that constitute the pseudobulk measurement.

**Figure S9.**
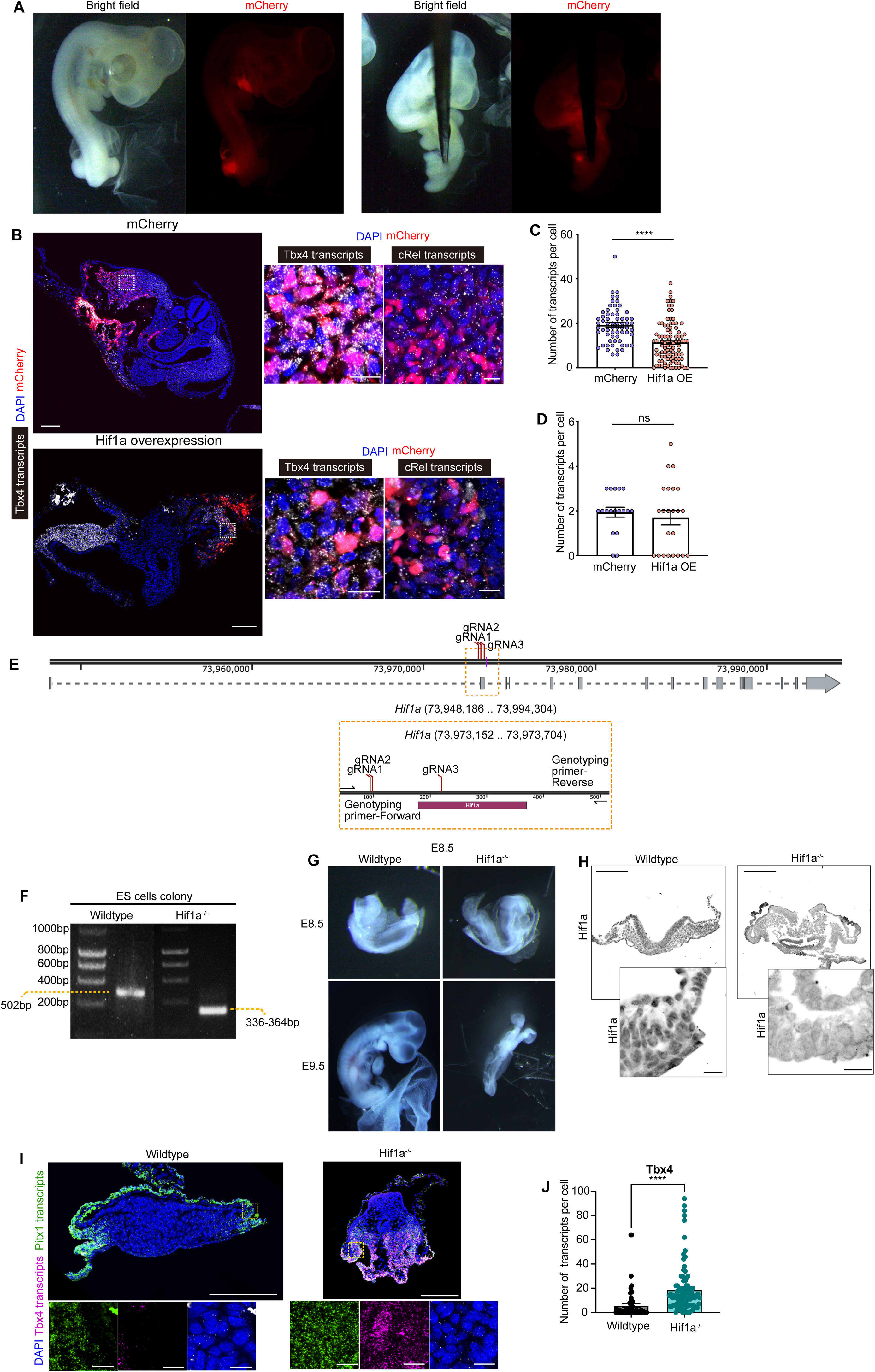
Hif1a expression does not regulate that of cRel in the mouse hindlimb field. (**A**) Stereoscope images of embryos overexpressing pCAG-mCherry (control) and pCAG-Hif1a+pCAG-mCherry for 48 hours. (**B**) Sections from embryos with hindlimb tissues overexpressing pCAG-mCherry (control) and pCAG-Hif1a+pCAG-mCherry for 48 hours were stained with mCherry, Tbx4 HCR and DAPI. Magnified images showed hindlimb cells. (**C**) Quantifications of Tbx4 transcripts in cells overexpressing pCAG-mCherry (control) and pCAG-Hif1a+pCAG-mCherry for 48 hours as in **B**. (**D**) Quantifications of cRel transcripts in cells overexpressing pCAG-mCherry (control) and pCAG-Hif1a+pCAG-mCherry for 48 hours as in **B**. (**E**) Genomic locus of Hif1a and positions of gRNA and genotyping primers. Three guide RNAs were used to digest out the fragment of DNA in Hif1a locus that contains the start coding sites. Genotyping primers were designed to amplify the sequences and select those that are homozygous mutants that do not contain the targeted DNA fragments (wildtype – 502bp; mutants – 336-364bp). (**F**) Genotyping of single ES cell colonies transfected with gRNAs and Cas9 protein. Right lane showed the clone used to generate Hif1a null embryos (selected from 48 colonies) whereas the left lane shows the wildtype control clones used for control experiments. (**G**) Stereoscope images of chimeric embryos carrying wildtype ES cells vs Hif1a null ES cells. Hif1a null embryos display morphological defects such as the failure of neural tube closure and the lack of growth. (**H**) Validation of Hif1a protein depletion by Hif1a immunostaining towards wildtype and Hif1a null embryos dissected at E8.5. (**I**) Wildtype and Hif1a null embryos generated from ES cells – embryo chimera approach were dissected at E8.5 and stained with Pitx1 HCR, Tbx4 HCR and DAPI. Magnified regions indicate hindlimb field cells. (**J**) Quantifications of Tbx4 transcripts on wildtype and Hif1a null embryos as shown in **I**. Each dots represents a quantified cell. N=2 embryos for whildtype and N=3 embryos for Hif1a null embryos. N=2 experiments. Scale bars for overview images, 150μm; scale bars for magnified images, 15μm.

**Fig. S10.**
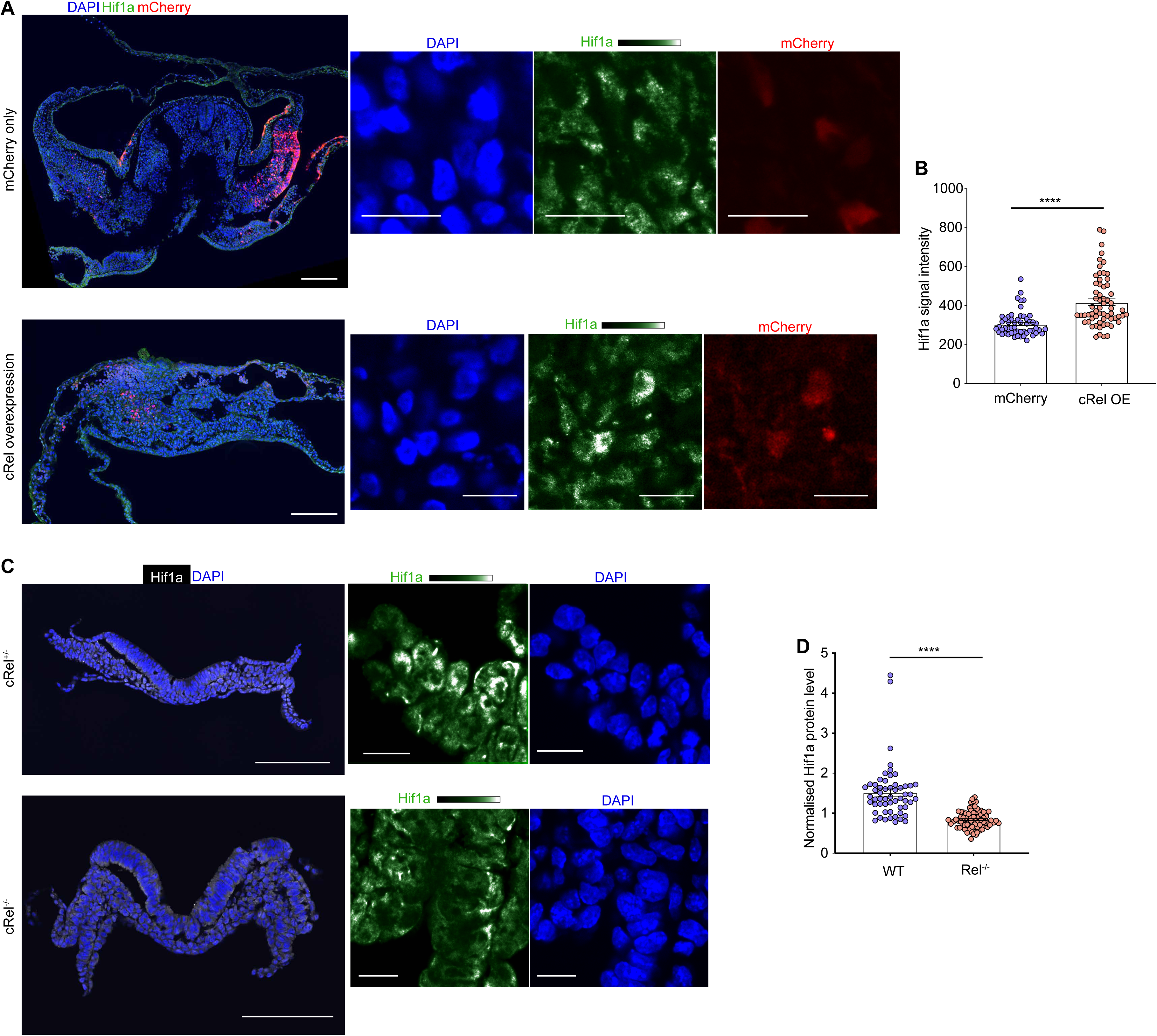
cRel promotes Hif1a’s expression in the mouse hindlimb field. (**A**) Representative images of chicken embryos electrporated with pCAG-mCherry (control) or pCAG-cRel:ZsGreen construct for 24 hours and immunostained with mCherry, Hif1a and DAPI. Magnified images represent the hindlimb tissues. (**B**) Quantifications of normalized Hif1a signal (nuclear against non-nuclear) in hindlimb cells from conditions shown in **A**. Each dot represents a quantified cell. N=2 experiments. ****p<0.0001, Mann Whitney test. (**C**) Representative images of cRel heterozygous or homozygous mouse embryos at E8.5 sectioned at the hindlimb level and stained with Hif1a and DAPI. (**D**) Quantifications of normalized Hif1a signal (nuclear against non-nuclear) in hindlimb field cells from conditions shown in **C**. Each dots represents a quantified cell. N=1 experiment. **** p<0.0001, Mann Whitney test.

## References

1. Richardson, M.K., Gobes, S.M., van Leeuwen, A.C., Polman, J.A., Pieau, C., and Sanchez-Villagra, M.R. (2009). Heterochrony in limb evolution: developmental mechanisms and natural selection. J Exp Zool B Mol Dev Evol 312, 639–664. 10.1002/jez.b.21250.

2. Weisbecker, V., Goswami, A., Wroe, S., and Sanchez-Villagra, M.R. (2008). Ossification heterochrony in the therian postcranial skeleton and the marsupial-placental dichotomy. Evolution 62, 2027–2041. 10.1111/j.1558-5646.2008.00424.x.

3. Keyte, A.L., and Smith, K.K. (2014). Heterochrony and developmental timing mechanisms: changing ontogenies in evolution. Semin Cell Dev Biol 34, 99–107. 10.1016/j.semcdb.2014.06.015.

4. Gros, J., and Tabin, C.J. (2014). Vertebrate limb bud formation is initiated by localized epithelial-to-mesenchymal transition. Science 343, 1253–1256. 10.1126/science.1248228.

5. Sekine, K., Ohuchi, H., Fujiwara, M., Yamasaki, M., Yoshizawa, T., Sato, T., Yagishita, N., Matsui, D., Koga, Y., Itoh, N., and Kato, S. (1999). Fgf10 is essential for limb and lung formation. Nat Genet 21, 138–141. 10.1038/5096.

6. Takeuchi, J.K., Koshiba-Takeuchi, K., Suzuki, T., Kamimura, M., Ogura, K., and Ogura, T. (2003). Tbx5 and Tbx4 trigger limb initiation through activation of the Wnt/Fgf signaling cascade. Development 130, 2729–2739. 10.1242/dev.00474.

7. Barrow, J.R., Thomas, K.R., Boussadia-Zahui, O., Moore, R., Kemler, R., Capecchi, M.R., and McMahon, A.P. (2003). Ectodermal Wnt3/beta-catenin signaling is required for the establishment and maintenance of the apical ectodermal ridge. Genes Dev 17, 394–409. 10.1101/gad.1044903.

8. Rallis, C., Bruneau, B.G., Del Buono, J., Seidman, C.E., Seidman, J.G., Nissim, S., Tabin, C.J., and Logan, M.P. (2003). Tbx5 is required for forelimb bud formation and continued outgrowth. Development 130, 2741–2751. 10.1242/dev.00473.

9. Naiche, L.A., and Papaioannou, V.E. (2003). Loss of Tbx4 blocks hindlimb development and affects vascularization and fusion of the allantois. Development 130, 2681–2693. 10.1242/dev.00504.

10. Kawakami, Y., Capdevila, J., Buscher, D., Itoh, T., Rodriguez Esteban, C., and Izpisua Belmonte, J.C. (2001). WNT signals control FGF-dependent limb initiation and AER induction in the chick embryo. Cell 104, 891–900. 10.1016/s0092-8674(01)00285-9.

11. Choi, H.M.T., Schwarzkopf, M., Fornace, M.E., Acharya, A., Artavanis, G., Stegmaier, J., Cunha, A., and Pierce, N.A. (2018). Third-generation in situ hybridization chain reaction: multiplexed, quantitative, sensitive, versatile, robust. Development 145. 10.1242/dev.165753.

12. Takeuchi, J.K., Koshiba-Takeuchi, K., Matsumoto, K., Vogel-Hopker, A., Naitoh-Matsuo, M., Ogura, K., Takahashi, N., Yasuda, K., and Ogura, T. (1999). Tbx5 and Tbx4 genes determine the wing/leg identity of limb buds. Nature 398, 810–814. 10.1038/19762.

13. Ohuchi, H., Nakagawa, T., Yamamoto, A., Araga, A., Ohata, T., Ishimaru, Y., Yoshioka, H., Kuwana, T., Nohno, T., Yamasaki, M., et al. (1997). The mesenchymal factor, FGF10, initiates and maintains the outgrowth of the chick limb bud through interaction with FGF8, an apical ectodermal factor. Development 124, 2235–2244. 10.1242/dev.124.11.2235.

14. Ng, J.K., Kawakami, Y., Buscher, D., Raya, A., Itoh, T., Koth, C.M., Rodriguez Esteban, C., Rodriguez-Leon, J., Garrity, D.M., Fishman, M.C., and Izpisua Belmonte, J.C. (2002). The limb identity gene Tbx5 promotes limb initiation by interacting with Wnt2b and Fgf10. Development 129, 5161–5170. 10.1242/dev.129.22.5161.

15. Rodriguez-Esteban, C., Tsukui, T., Yonei, S., Magallon, J., Tamura, K., and Izpisua Belmonte, J.C. (1999). The T-box genes Tbx4 and Tbx5 regulate limb outgrowth and identity. Nature 398, 814–818. 10.1038/19769.

16. Duboule, D. (1994). Temporal colinearity and the phylotypic progression: a basis for the stability of a vertebrate Bauplan and the evolution of morphologies through heterochrony. Dev Suppl, 135-142.

17. Minguillon, C., Nishimoto, S., Wood, S., Vendrell, E., Gibson-Brown, J.J., and Logan, M.P. (2012). Hox genes regulate the onset of Tbx5 expression in the forelimb. Development 139, 3180–3188. 10.1242/dev.084814.

18. Moreau, C., Caldarelli, P., Rocancourt, D., Roussel, J., Denans, N., Pourquie, O., and Gros, J. (2019). Timed Collinear Activation of Hox Genes during Gastrulation Controls the Avian Forelimb Position. Curr Biol 29, 35–50 e34. 10.1016/j.cub.2018.11.009.

19. Tanaka, M., Jokubaitis, V., Wood, C., Wang, Y., Brouard, N., Pera, M., Hearn, M., Simmons, P., and Nakayama, N. (2009). BMP inhibition stimulates WNT-dependent generation of chondrogenic mesoderm from embryonic stem cells. Stem Cell Res 3, 126–141. 10.1016/j.scr.2009.07.001.

20. Mori, S., Sakakura, E., Tsunekawa, Y., Hagiwara, M., Suzuki, T., and Eiraku, M. (2019). Self-organized formation of developing appendages from murine pluripotent stem cells. Nat Commun 10, 3802. 10.1038/s41467-019-11702-y.

21. Logan, M., and Tabin, C.J. (1999). Role of Pitx1 upstream of Tbx4 in specification of hindlimb identity. Science 283, 1736–1739. 10.1126/science.283.5408.1736.

22. Duboc, V., and Logan, M.P. (2011). Pitx1 is necessary for normal initiation of hindlimb outgrowth through regulation of Tbx4 expression and shapes hindlimb morphologies via targeted growth control. Development 138, 5301–5309. 10.1242/dev.074153.

23. Itou, J., Kawakami, H., Quach, T., Osterwalder, M., Evans, S.M., Zeller, R., and Kawakami, Y. (2012). Islet1 regulates establishment of the posterior hindlimb field upstream of the Hand2-Shh morphoregulatory gene network in mouse embryos. Development 139, 1620–1629. 10.1242/dev.073056.

24. Duboc, V., Sulaiman, F.A., Feneck, E., Kucharska, A., Bell, D., Holder-Espinasse, M., and Logan, M.P.O. (2021). Tbx4 function during hindlimb development reveals a mechanism that explains the origins of proximal limb defects. Development 148. 10.1242/dev.199580.

25. Szeto, D.P., Rodriguez-Esteban, C., Ryan, A.K., O’Connell, S.M., Liu, F., Kioussi, C., Gleiberman, A.S., Izpisua-Belmonte, J.C., and Rosenfeld, M.G. (1999). Role of the Bicoid-related homeodomain factor Pitx1 in specifying hindlimb morphogenesis and pituitary development. Genes Dev 13, 484–494. 10.1101/gad.13.4.484.

26. Narkis, G., Tzchori, I., Cohen, T., Holtz, A., Wier, E., and Westphal, H. (2012). Isl1 and Ldb co-regulators of transcription are essential early determinants of mouse limb development. Dev Dyn 241, 787–791. 10.1002/dvdy.23761.

27. Ferrer-Vaquer, A., Piliszek, A., Tian, G., Aho, R.J., Dufort, D., and Hadjantonakis, A.K. (2010). A sensitive and bright single-cell resolution live imaging reporter of Wnt/ss-catenin signaling in the mouse. BMC Dev Biol 10, 121. 10.1186/1471-213X-10-121.

28. Takemoto, T., Abe, T., Kiyonari, H., Nakao, K., Furuta, Y., Suzuki, H., Takada, S., Fujimori, T., and Kondoh, H. (2016). R26-WntVis reporter mice showing graded response to Wnt signal levels. Genes Cells 21, 661–669. 10.1111/gtc.12364.

29. Molotkov, A., Molotkova, N., and Duester, G. (2005). Retinoic acid generated by Raldh2 in mesoderm is required for mouse dorsal endodermal pancreas development. Dev Dyn 232, 950–957. 10.1002/dvdy.20256.

30. Cunningham, T.J., Zhao, X., Sandell, L.L., Evans, S.M., Trainor, P.A., and Duester, G. (2013). Antagonism between retinoic acid and fibroblast growth factor signaling during limb development. Cell Rep 3, 1503–1511. 10.1016/j.celrep.2013.03.036.

31. Jeong, S., Rebeiz, M., Andolfatto, P., Werner, T., True, J., and Carroll, S.B. (2008). The evolution of gene regulation underlies a morphological difference between two Drosophila sister species. Cell 132, 783–793. 10.1016/j.cell.2008.01.014.

32. Menke, D.B., Guenther, C., and Kingsley, D.M. (2008). Dual hindlimb control elements in the Tbx4 gene and region-specific control of bone size in vertebrate limbs. Development 135, 2543–2553. 10.1242/dev.017384.

33. Young, J.J., Grayson, P., Edwards, S.V., and Tabin, C.J. (2019). Attenuated Fgf Signaling Underlies the Forelimb Heterochrony in the Emu Dromaius novaehollandiae. Curr Biol 29, 3681–3691 e3685. 10.1016/j.cub.2019.09.014.

34. Alerasool, N., Segal, D., Lee, H., and Taipale, M. (2020). An efficient KRAB domain for CRISPRi applications in human cells. Nat Methods 17, 1093–1096. 10.1038/s41592-020-0966-x.

35. Mic, F.A., Haselbeck, R.J., Cuenca, A.E., and Duester, G. (2002). Novel retinoic acid generating activities in the neural tube and heart identified by conditional rescue of Raldh2 null mutant mice. Development 129, 2271–2282. 10.1242/dev.129.9.2271.

36. Savory, J.G., Bouchard, N., Pierre, V., Rijli, F.M., De Repentigny, Y., Kothary, R., and Lohnes, D. (2009). Cdx2 regulation of posterior development through non-Hox targets. Development 136, 4099–4110. 10.1242/dev.041582.

37. Bell, C.C., Amaral, P.P., Kalsbeek, A., Magor, G.W., Gillinder, K.R., Tangermann, P., di Lisio, L., Cheetham, S.W., Gruhl, F., Frith, J., et al. (2016). The Evx1/Evx1as gene locus regulates anterior-posterior patterning during gastrulation. Sci Rep 6, 26657. 10.1038/srep26657.

38. Qiu, C., Martin, B.K., Welsh, I.C., Daza, R.M., Le, T.M., Huang, X., Nichols, E.K., Taylor, M.L., Fulton, O., O’Day, D.R., et al. (2024). A single-cell time-lapse of mouse prenatal development from gastrula to birth. Nature 626, 1084–1093. 10.1038/s41586-024-07069-w.

39. Yin, K., and Juurlink, B.H. (2000). Regulation of expression of nuclear factor kappa B RelA in oligodendrocytes: effect of hypoxia. Neuroreport 11, 1877–1881. 10.1097/00001756-200006260-00015.

40. Jiang, Y., Zhu, Y., Wang, X., Gong, J., Hu, C., Guo, B., Zhu, B., and Li, Y. (2015). Temporal regulation of HIF-1 and NF-kappaB in hypoxic hepatocarcinoma cells. Oncotarget 6, 9409–9419. 10.18632/oncotarget.3352.

41. Carroll, C., Engström, N., Nilsson, P.F., Haxen, E.R., Mohlin, S., Berg, P., Glud, R.N., and Hammarlund, E.U. (2021). Hypoxia Generated by Avian Embryo Growth Induces the HIF-α Response and Critical Vascularization. Frontiers in Ecology and Evolution 9. 10.3389/fevo.2021.675800.

42. Aguilera-Castrejon, A., Oldak, B., Shani, T., Ghanem, N., Itzkovich, C., Slomovich, S., Tarazi, S., Bayerl, J., Chugaeva, V., Ayyash, M., et al. (2021). Ex utero mouse embryogenesis from pre-gastrulation to late organogenesis. Nature 593, 119–124. 10.1038/s41586-021-03416-3.

43. Sakai, D., Murakami, Y., Shigeta, D., Tomosugi, M., Sakata-Haga, H., Hatta, T., and Shoji, H. (2023). Glycolytic activity is required for the onset of neural plate folding during neural tube closure in mouse embryos. Frontiers in Cell and Developmental Biology 11. 10.3389/fcell.2023.1212375.

44. Culshaw, L.H., Savery, D., Greene, N.D.E., and Copp, A.J. (2019). Mouse whole embryo culture: Evaluating the requirement for rat serum as culture medium. Birth Defects Res 111, 1165–1177. 10.1002/bdr2.1538.

45. Linde-Medina, M., and Smit, T.H. (2021). Molecular and Mechanical Cues for Somite Periodicity. Front Cell Dev Biol 9, 753446. 10.3389/fcell.2021.753446.

46. Pijuan-Sala, B., Griffiths, J.A., Guibentif, C., Hiscock, T.W., Jawaid, W., Calero-Nieto, F.J., Mulas, C., Ibarra-Soria, X., Tyser, R.C.V., Ho, D.L.L., et al. (2019). A single-cell molecular map of mouse gastrulation and early organogenesis. Nature 566, 490–495. 10.1038/s41586-019-0933-9.

47. Iyer, N.V., Kotch, L.E., Agani, F., Leung, S.W., Laughner, E., Wenger, R.H., Gassmann, M., Gearhart, J.D., Lawler, A.M., Yu, A.Y., and Semenza, G.L. (1998). Cellular and developmental control of O2 homeostasis by hypoxia-inducible factor 1 alpha. Genes Dev 12, 149–162. 10.1101/gad.12.2.149.

48. Keyte, A.L., and Smith, K.K. (2010). Developmental origins of precocial forelimbs in marsupial neonates. Development 137, 4283–4294. 10.1242/dev.049445.

49. Harrison, L.B., and Larsson, H.C. (2008). Estimating evolution of temporal sequence changes: a practical approach to inferring ancestral developmental sequences and sequence heterochrony. Syst Biol 57, 378–387. 10.1080/10635150802164421.

50. Lyne, A. (1964). Observations on the breeding and growth of the marsupial Perameles nasuta Geoffroy, with notes on other bandicoots. Australian Journal of Zoology 12, 322–339.

51. Chew, K.Y., Shaw, G., Yu, H., Pask, A.J., and Renfree, M.B. (2014). Heterochrony in the regulation of the developing marsupial limb. Dev Dyn 243, 324–338. 10.1002/dvdy.24062.

52. de Jesus, T.J., and Ramakrishnan, P. (2020). NF-kappaB c-Rel Dictates the Inflammatory Threshold by Acting as a Transcriptional Repressor. iScience 23, 100876. 10.1016/j.isci.2020.100876.

53. Kontgen, F., Grumont, R.J., Strasser, A., Metcalf, D., Li, R., Tarlinton, D., and Gerondakis, S. (1995). Mice lacking the c-rel proto-oncogene exhibit defects in lymphocyte proliferation, humoral immunity, and interleukin-2 expression. Genes Dev 9, 1965–1977. 10.1101/gad.9.16.1965.

54. Nerurkar, N.L., Lee, C., Mahadevan, L., and Tabin, C.J. (2019). Molecular control of macroscopic forces drives formation of the vertebrate hindgut. Nature 565, 480–484. 10.1038/s41586-018-0865-9.

55. Hao, Y., Stuart, T., Kowalski, M.H., Choudhary, S., Hoffman, P., Hartman, A., Srivastava, A., Molla, G., Madad, S., Fernandez-Granda, C., and Satija, R. (2024). Dictionary learning for integrative, multimodal and scalable single-cell analysis. Nat Biotechnol 42, 293–304. 10.1038/s41587-023-01767-y.

